# Impact of thermal variation on the transcriptome of *Plasmodium falciparum*- infected *Anopheles stephensi*

**DOI:** 10.64898/2026.07.17.739141

**Authors:** Ashutosh K Pathak, Shannon Quek, Ritu Sharma, Justine C Shiau, Matthew B. Thomas, Grant L. Hughes, Courtney C Murdock

**Author notes:** equal contribution (Ashutosh K Pathak, Shannon Quek).

## Abstract

Temperature is a key determinant of malaria transmission, influencing both parasite development and mosquito physiology, yet the underlying mechanisms remain poorly understood. Here, we examined how temperature and time modulate gene expression in *Anopheles stephensi* infected with *Plasmodium falciparum*. Using RNA-sequencing over 1-19 days post-blood meal and three temperature regimes (20, 24, and 28°C with diurnal fluctuations of 9°C), we characterize transcriptome responses to infection with *P. falciparum* at the site of infection in the midgut, and systemically, in the carcasses. Oocyst prevalence and density declined over the thermal gradient, albeit with distinct, non-linear temporal dynamics in parasite development rates. Although infection contributed minimally to global variation in gene expression relative to temperature and time, infection-associated genes in the midgut showed coordinated transcriptional responses enriched in canonical *Plasmodium* associated extracellular, proteolytic, immune, and metabolic functions; notably, decline in oocyst infections in the midguts over the thermal gradient was reflected in reduced expression of immune genes known to regulate *P. falciparum*. Network analysis demonstrated that these genes participate in a significantly interconnected protein–protein interaction network, within which a small number of high betweenness centrality proteins act as bottlenecks linking immune, metabolic, reproductive and behavioral processes. Our results suggest responses to infection may be mediated through coordinated physiological networks rather than large-scale transcriptional changes. Our findings also indicate that differences in thermal conditions may be an important factor when comparing mechanisms of vector– parasite interactions between *Plasmodium* species. Together, our results highlight the importance of integrating thermal context into mechanistic studies of vector–parasite interactions.

## Introduction

When the initial goal to achieve malaria elimination by 2030 was set forth in 2000, it was considered ambitious primarily due to logistical challenges of implementation on the ground ^1^. In addition to these challenges, global landscapes and climate are changing resulting in shifts in the distribution, abundance, and seasonality of mosquito populations and malaria cases ^2^ . One of the fasting growing landscapes on the planet are urban environments, which are also experiencing intensifying heat extreme events due to our warming climate. Initially, urbanization was predicted to work synergistically with rural control efforts in reducing malaria burdens due to a general lack of suitable habitat for native Anopheline vectors. Further, heat extreme events could further decrease suitability for parasite transmission ^3–6^. However, native Anopheline vectors are adapting to rural environments due to peri-urban agriculture, and the emergence of the invasive S. Asian urban malaria vector, *Anopheles stephensi*, have resulted in increasing incidence of *P. falciparum* ^4^ in urban environments. Its re-appearance in areas that eliminated autochthonous transmission by indigenous *Anopheles* populations for at least two decades is a reminder of the fragility of gains achieved over the past two decades. As *An. stephensi* has gained a strong foothold in regions of East Africa and is spreading across central and Western Africa, understanding how the mosquito adapts to its new environment, and how this modifies vectorial capacity may be crucial to understanding and managing disease transmission ^7,8^.

Parasite transmission and disease incidence are dependent on how the vector responds to multiple environmental factors that determine the timing, persistence, and overall magnitude of malaria cases that occur ^1,9,2^. Variation in the thermal environment is one of the best-described regulators of vectorial capacity. Metabolic theory of ecology suggests that beyond a minimum critical temperature (*CT_min_*), the rate and efficiency of basic biological processes increase as they approach an intermediate temperature that maximizes performance (thermal optimum: *T_opt_*). As temperatures warm above the *T_opt_*, the efficiency of biological reactions decline rapidly until organismal death occurs at some maximum critical temperature (*CT_max_*) ^6,10–13^. Furthermore, temperature is also known to affect parasite survival and establishment ^13^ within the mosquito vector, though it is unclear if this occurs via a direct effect of temperature on parasite establishment, replication, and dissemination, or an indirect effect on parasite fitness through changes in mosquito immune-physiology^14,15^. It is known that the parasite is particularly susceptible to thermal variation in the first few hours post-ingestion^16–18^, when mating between male and female gametes in the lumen of the mosquito midgut is followed by differentiation of the zygote into ookinetes. Once the ookinetes have traversed the epithelial lining of the midgut and established in the basal lamina, oocysts are relatively less susceptible to variation in the external environment ^16–19^. This protection is lost as sporozoites leave the oocyst to begin their migration to the salivary glands ^20–22^. Studies have attempted to define the effects of temperature on mosquito and parasite traits that drive transmission by placing mosquitoes across varying thermal conditions and measuring the performance and variation in performance of a given phenotype ^6,10–13^ . However, less attention has been dedicated toward understanding the effects of temperature on the mechanisms driving trait variation across temperature or the outcome of mosquito-parasite interactions ^13,23,24^, which is necessary to predict physiological trade-offs and evolutionary outcomes in response to environmental change ^25,26^.

To better understand how temperature may affect mosquito gene expression and life history traits over time, we previously reported transcriptomes of *An. stephensi* housed under daily mean temperatures of 20, 24 and 28°C, with diurnal fluctuations of 9°C (+5°C / –4°C) around each mean (20 DTR 9°C, 24 DTR 9°C and 28 DTR 9°C respectively herein) ^27^. The midgut and carcass transcriptomes were resolved from 1-19 days post-blood meal and highlighted the trade-offs associated with blood feeding behavior, oogenesis, oviposition and ageing across different temperatures without the impact of *Plasmodium* infection. In this study, we develop this further by looking at the effect of both temperature and *P. falciparum* infection on temporal gene expression in the midguts and carcasses.

## Methods

### Study design

The study design and temperature regimens were identical to our previous study^27^ (Figure 1). Two days before the blood meal, 750, 3–5-day old female, host-seeking *An. stephensi* were aspirated into six polyester cages (32.5 cm^3^, BugDorm, MegaView Science, Taiwan). Two cages each were transferred to environmental chambers (Percival Scientific, Perry, IA) maintained at 20 DTR 9°C, 24 DTR 9°C and 28 DTR 9°C. After mosquitoes had been acclimated to the three temperatures for 48h, at ZT18h, mosquitoes in one cage from each chamber was offered a meal of naïve RBCs and serum suspension (“controls”), while the second cage was offered the same blood meal but spiked with *P. falciparum* NF54 propagated in primary RBCs in vitro (0.3% gametocytemia) ^22,28^. Mosquitoes were allowed to feed for 15 min after which 500, fully engorged mosquitoes were then returned to the respective temperature regime.

**Figure 1:**
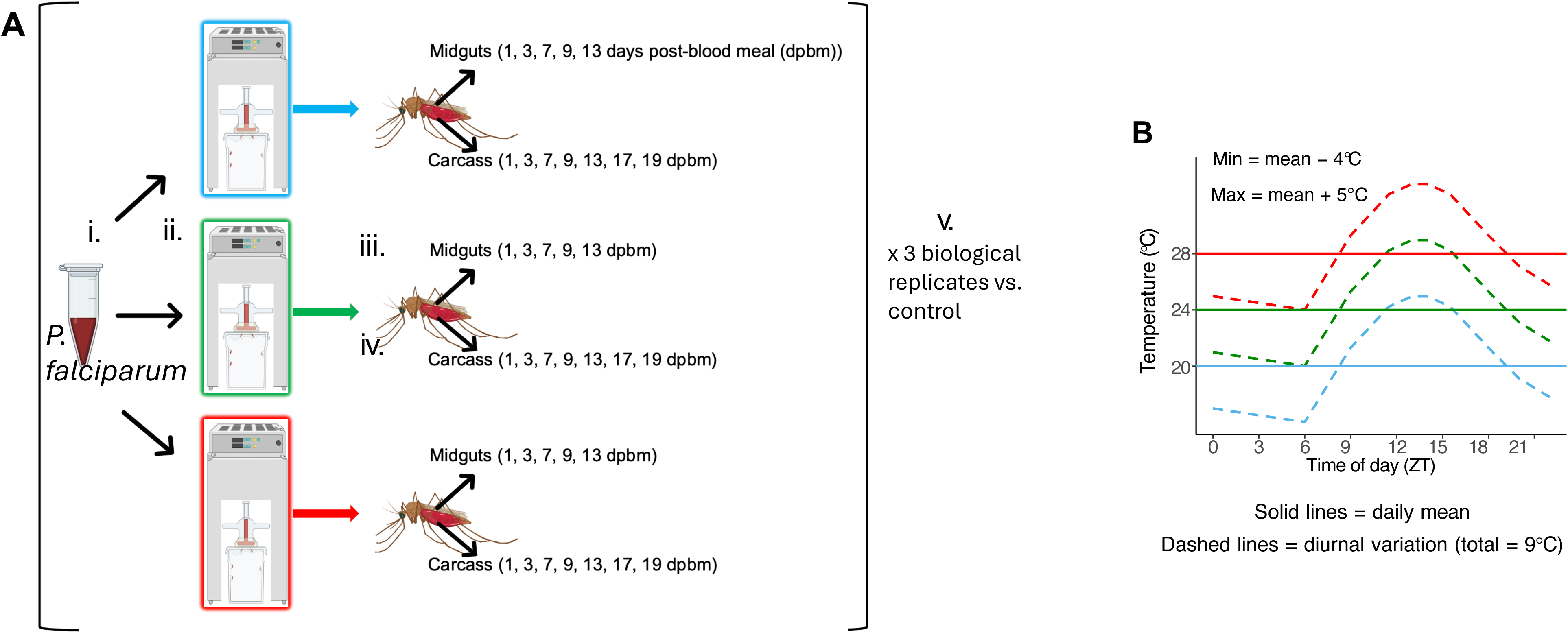
Graphical overview of experimental design.

At ZT15 h on 1, 3, 7, 9, 13, 17, and 19 days post-blood meal (dpbm), approximately 20–36 mosquitoes were sampled from control and *P. falciparum*-exposed cages at each of the three temperatures. For exposed groups, mosquitoes were screened with the goal of obtaining 10 individuals with microscopically confirmed oocyst-positive midguts when parasite development was sufficiently advanced for infection status to be verified (see next paragraph for details). At time points when oocysts could not be reliably confirmed, pools were generated from parasite-exposed mosquitoes of unknown infection status. To quantify transcriptional dynamics at the local site of infection, midguts from 10 mosquitoes per treatment group were pooled at 1, 3, 7, 9, and 13 dpbm. The remaining tissues from the same individuals were pooled separately as carcasses to assess systemic transcriptional dynamics; carcasses were collected at all seven time points. This sampling design was repeated across three biological replicates. In each replicate, RNA-seq was performed on 72 samples: 30 midgut pools [5 time points × 2 exposure statuses × 3 temperatures] and 42 carcass pools [7 time points × 2 exposure statuses × 3 temperatures], comprising 36 control and 36 parasite-exposed samples.

For data analysis, our approach considered two potential (albeit unavoidable) limitations to our study design. First, although our goal was to characterize transcriptomes from mosquitoes that were known to be infected (i.e., oocyst-positive), temperature-dependent differences in parasite development rates meant infection status could not be confirmed at 1 and 3 dpbm at any temperature, and from 9 dpbm onwards at the coolest temperature (20°C DTR 9°C). As such, a subset of samples (7 per replicate; 21 total) represents parasite-exposed mosquitoes of unknown infection status, whereas the remaining samples (14 per replicate; 42 total) were derived from individuals with confirmed infections. We account for this in our statistical analyses of the transcriptomes by identifying genes whose expression is affected by all three treatments of temperature, time and infection status. Second, elevated temperatures are associated with reduced mosquito survival ^10^ and exacerbated by infection and parasite burden ^29^; in other words, depending on the temperature, mosquitoes collected later in infection may disproportionately represent individuals with lower parasite densities, or mosquitoes that successfully cleared infection early on. If true, transcriptomes would be biased towards different sub-populations depending on time of sampling and potentially exacerbated by temperature. To mitigate this bias, we monitored infection dynamics (oocyst prevalence and density) at 7, 9, 13, 17, and 19 dpbm, as well as at intermediate time points (5, 11, and 15 dpbm). While these additional time points were not included in RNAseq, they provided a higher-resolution view of infection dynamics across 1–19 dpbm and allowed us to detect any major temperature-dependent shifts in parasite development, which in turn could be indicative, albeit indirect of differences in survival; moreover, to increase the sensitivity of detecting any changes, and the fact that dynamics are inherently non-linear and may differ in both shape and magnitude across temperatures, parasite infections were analyzed using generalized additive mixed models (GAMMs).

Finally, although our initial goal was to characterize the transcriptomes in the salivary glands of these *P. falciparum*-infected mosquitoes, RNA yields were too low for any meaningful analyses. While the relatively small size of this organ may have required a significantly large number of glands to be pooled for robust analyses, the strong reduction in prevalence at 28 DTR 9°C compared to the two cooler temperatures ^22^ would also make it difficult to distinguish between *P. falciparum*-specific and general temperature driven gene expression profiles. To demonstrate the effect of temperature on sporozoite prevalence, we analyzed data from seven independent infections where sporozoite prevalence was measured in the salivary glands.

### Parasite cultures

Unless stated otherwise, all biological and chemical reagents and consumables were purchased from Thermo-Fisher Scientific (Waltham, MA). *P. falciparum* NF54 (obtained through BEI Resources, NIAID, NIH: *Plasmodium falciparum*, Strain NF54 (Patient Line E), MRA-1000, contributed by Megan G. Dowler) were cultured with O^+^ cryogenically preserved RBCs in vitro (Valley Biomedical, Winchester, VA) ^27,28^. Asexual cultures of the parasites were maintained in 5% RBCs resuspended in RPMI-1640 media supplemented with HEPES, sodium bicarbonate, hypoxanthine and 10% human serum (A^+^) (complete media). Parasitemia was monitored by Giemsa staining every 48-72h. Once asexual cultures had reached 3-5% parasitemia, gametocytes cultures were initiated in T75 flasks at a starting density of 0.5- 1% in complete media and freshly thawed cryogenically preserved RBCs at a 5% hematocrit. Rates of gametocytogenesis were monitored at days 1, 4, 7 and 12 post-seeding. Assays for maturation of male gametocytes (exflagellation) were started at day 12, and once exflagellation centers were observed (usually 12-14 days post-seeding), cultures were used to prepare the infectious blood meals within 48h (usually 14-16 days post-seeding).

### Mosquito husbandry

Our colony of *An. stephensi* was initiated from eggs kindly provided by the Walter Reed Army Institute of Research ca. 2015 ^27,28^. This is a wild-type strain classified as strain ‘Indian’, type ‘urban’. Colonies were propagated on a weekly basis with eggs oviposited by mosquitoes offered packed, O+ human RBCs supplemented with freshly thawed A+ human serum (Valley Biomedical, Winchester, VA). Larvae were propagated on a progressive feeding schedule of Hikari Cichlid Gold medium pellets (Hikari Inc, Hayward, CA), while adults were routinely maintained on 5% dextrose (w/v) and 0.05% PABA (w/v) as described previously ^28^.

### Infecting An. stephensi with P. falciparum

Two days (48h) before the infections, female *An. stephensi* (strain ‘Indian’, type ‘urban’), 3-5 days post- eclosion showing active host-seeking behavior were sorted into 32.5cm^3^ polyester mesh cages (Bugdorm Inc, Taiwan). Cages were acclimated in reach-in environmental chambers (Percival Scientific, Perry, IA) maintained at 20 DTR 9°C, 24 DTR 9°C and 28 DTR 9°C. On the day of infection, infectious blood meals were prepared from the parasite cultures by first concentrating the RBCs with centrifugation at 1,800g for 3 minutes at room temperature and half the maximum deceleration setting. These infected RBCs were then resuspended in freshly thawed cryogenically preserved RBCs and human serum to achieve a mature gametocyte density of ∼0.3% in the infectious blood meal and final hematocrit of 50% (1:1 RBC to serum ratio); control cages were offered the same cryogenically preserved RBCs diluted to a hematocrit of ∼50% with human serum ^27,28^. Both blood meals were aliquoted into water-jacked glass feeders lined with parafilm and maintained at 38°C and offered to the mosquitoes at ∼1800ZT. After 15 minutes, blood-engorged mosquitoes were returned to their respective chambers. Infectivity of the blood meal was confirmed by assessing male gametogenesis (or activation of male gametocytes).

### Collection of tissues from infected and control mosquitoes for RNAseq

At the indicated time points, ∼20–25 mosquitoes from each cage were directly aspirated into 70% ethanol, with midguts and carcasses from 10 individuals pooled into 50 μL and 500 μL of RNAlater respectively. As stated earlier, while it was not possible to confirm infection status at 1 and 3 dpbm as the oocysts were not visible, all tissues collected thereafter were from mosquitoes with ≥1oocyst in the midgut. Oocyst quantifications in the midguts were performed at 400× magnification with a Leica DM2500 under DIC optics ^22,28^. However, to avoid rupturing the oocysts with the coverslip, midguts from each individual were resuspended in a 10µl volume of chilled PBS in each well of a 6-well Polytetrafluoroethylene (PTFE) printed slides; the PTFE coating around each well prevents the coverslip from crushing the midguts while remaining thin enough to allow visualization with a microscope. After overnight storage in RNAlater at 2– 4 °C, samples were transferred to −80 °C until RNA purification and sequencing.

### RNA purification and sequencing

Total RNA was purified from the midguts and carcasses with a commercially available kit (PureLink RNA Mini Kit) as suggested by the manufacturer (Thermo-Fisher Scientific, Waltham, MA). RNA sequencing (RNAseq herein) was performed at the Georgia Genomics and Bioinformatics Core (GGBC) at the University of Georgia (Athens, GA). Concentration and quality checks for integrity of total RNA were assessed with a Bioanalyzer (Thermo-Fisher Scientific, Waltham, MA). Libraries were prepared from each sample with a KAPA stranded mRNA-seq kit as recommended by the manufacturer (Roche, Indianapolis, IN); mRNA was selected with oligo-dT beads and fragmented prior to generating cDNA using random hexamer priming. The quality and quantity of each library were estimated with a qubit or plate reader before pooling for RNAseq. Pooled libraries were run on a NextSeq 550 (Illumina Inc., San Diego, CA). De-multiplexing and trimming of adapter and barcode sequences were performed on BaseSpace (Illumina Inc., San Diego, CA).

### Data analyses and modeling

Parasite infections in the mosquitoes were modeled with GAMMs with the package “mgcv” (version 1.9- 3) ^30,31^ in RStudio (version 2025.05.0) running over the R open source software (version 4.5.0). Time was modeled as a smooth term with temperature as a categorical factor, with an interaction between two to allow temperature-specific temporal trajectories. Replicate was included as a random effect. In general, the effect of a smoothed predictor is expressed as effective degrees of freedom (EDF) ^31^. While EDF values ≤ 1 indicate a linear relationship with the dependent variable, EDFs > 1 indicate non-linearity, with larger values indicating greater non-linearity/curvature; for instance, an EDF of 2 indicates a quadratic/humped relationship.

The proportion of oocyst-positive mosquitoes (oocyst prevalence) was modeled with a binomial distribution (‘logit’ link) and oocyst densities with a type 2 negative binomial (‘log’ link). The significance of each term in the model was based on the extent (chi-squared, χ2 herein) to which it accounted for the unexplained deviance (or variation) in the dependent variable. Model diagnostics were performed with “gratia” package (version 0.10.0) ^32^. Model performance and selection was based on Akaike’s information theoretic criteria with the “bbmle” package (version 1.0.25.1) ^33^. All post hoc estimations and comparisons of slopes predicted by the model performed with the “emmeans” package, after adjusting for family wise error rates with the Bonferroni-Holm method (version 1.11.2) ^34^. Finally, the proportion of sporozoite- positive salivary glands were modeled with a betabinomial distribution with the package “glmmTMB” (version 1.1.14) ^35,36^, with temperature treated as a categorical predictor, with means at each temperature allowed to vary between the replicates. Model diagnostics were performed with the package “DHARMa” (version 0.5.0) ^37^.

### Differential expression analysis

Following the protocol of our previous publication, sequences were aligned to the *Anopheles stephensi* “Indian” strain reference genome (genome version GCA_000300775.2, annotation version AsteI2.3) using the program Hisat2 (v2.1.0, ). FeatureCounts (v2.0.1, ^38^) was used to obtain raw read count data from the sequencing files, and the resulting counts table used as input into the program DESeq2 (v1.49.1, ^39^) in R (v4.5.0, ^40^). No normalization or pre-filtering of genes and their counts was performed, as DESeq2 automatically performs both steps - first the normalization, then subsequent filtering of genes based on the mean of normalized counts as a filter statistic. This is one of the recommendations by DESeq2’s designers (see vignette). Normalized gene counts were used to generate a PCA plot and sample-to- sample distance matrices in DESeq2. This was performed twice, either keeping counts from carcasses and midguts separate, or combining them together into a large dataset. This was done for quality control and identification of sample clustering, and to identify any outliers. Five samples appeared as distinct outliers (no clustering with any biological replicate or any other sample) and were thus excluded from analysis. All experimental conditions had a minimum of two biological replicates, even after accounting for outliers or poorly sequenced samples.

To investigate the impact of infection, time since bloodmeal (days 1-19, depending on tissue being analyzed), and temperature (20°C, 24°C and 28°C, DTR 9°C) on gene expression in mosquitoes across three different temperatures, we utilized DESeq2’s built-in likelihood ratio test (LRT) analysis functionality. To more appropriately model the tripartite effects in the system, we implemented a full interaction model within the DESeq2 framework with all three experimental factors (full model of ∼ time * temperature * infection). The LRT analysis then utilizes a ‘reduced’ model, where one term of the full model is removed from the analysis. This effectively tests if the removed term accounts for significant differential expression observed in the genes (e.g., a reduced model of ∼ time * temperature, tests for the role of Infection in contributing to differential expression). The goal of this likelihood ratio test (LRT) framework is to identify genes whose expression is consistently influenced by infection across all experimental conditions. Although this approach is more conservative than, for instance, pairwise comparisons between infected and control mosquitoes at individual time points or temperatures, it reduces the likelihood of identifying condition-specific or transient expression changes as infection-associated effects. Importantly, the factorial model also accounts for the non-independence of observations across time points and temperature treatments.

Genes were considered differentially expressed at a false discovery rate < 0.05 (FDR, estimated as Benjamini–Hochberg p-values adjusted for multiple testing). However, no fold-change metric can be calculated directly as the analysis considers multiple comparisons across the time-series experiment. Overlaps in gene lists between the models were visualized with Upset plots using the R package ComplexUpset (v1.3.3, ^41,42^) and queried using ComplexHeatmap (v2.26.0, ^43^). Further analysis was focused on genes that were statistically significant based on an overlap between infection, time, and temperature, as we had already investigated genes that were statistically significant based on the other two factors in a previous publication ^27^. These gene lists were used to perform direct comparisons of gene expression between infected and uninfected tissues at the same time-point and rearing temperature. Fold-changes were then extracted and plotted onto a heatmap in R using pheatmap (v1.0.13, ^45^), and the heatmap dendrogram was used to identify clusters of genes that showed co-regulation of expression.

### Identification of gene expression profiles

This clustering analysis allows us to group genes together based on similar expression profiles across the time-series and can give us insights into how these gene groups, or even whole biochemical pathways are responding across the different studied temperatures. To illustrate how the genes may migrate between clusters based on rearing temperature, we generated Upset plots using the R package ComplexUpset ^41,42^, using gene lists extracted from each cluster. These genes were further analyzed using the STRING database to identify enriched functional categories and protein–protein interaction (PPI) networks. Enrichment analysis identifies overrepresented biological functions based on gene counts within categories, whereas network analysis identifies genes that are functionally connected through protein–protein interactions. As these approaches capture distinct aspects of the data, the resulting gene sets may not fully overlap.

Enrichment analysis and protein–protein interaction networks were constructed using the STRING database (v12.0) via the stringApp (v2.2.0; released December 2024) implemented in Cytoscape (v3.10.4) ^46^; with default thresholds used for both. The resulting PPI network was used to identify proteins that bridge multiple biological processes (‘bottlenecks’) based on betweenness centrality (BC), a measure of how frequently a node lies on the shortest paths between other nodes ^47^. Nodes with high BC are therefore positioned to connect otherwise separate functional modules, independent of their expression magnitude in any single condition. Finally, Markov Cluster Algorithm (MCL) was used to identify modules within the network, with genes in each module subjected to enrichment analysis to determine functional associations (FDR < 0.05); categories/terms with the lowest FDR values were then annotated onto the network.

## Results

### Effect of temperature on oocyst prevalence and densities

We found significant effects of temperature on mean oocyst prevalence and oocyst densities per midgut. Mean oocyst prevalence was lowest in replicate 1 (27.7 ± 3.96%) but higher in replicates 2 and 3 (58.4 ± 3.79% and 64.6 ± 3.49%, respectively; Supplementary figure 1). After accounting for this substantial variation between replicates in the model (χ^2^ = 154.6, p<0.001), we found significant declines in mean oocyst prevalence with warming temperature (χ^2^ = 85.9, p<0.001), with lowest prevalence observed at 28 DTR 9°C relative to 20 DTR 9°C (20 DTR 9°C vs. 28 DTR 9°C: β = −1.39, p < 0.001) and intermediate prevalence at 24 DTR 9°C (20 DTR 9°C vs. 24 DTR 9°C: β = −0.36, p = 0.037) (Figure 2A). While the overall smooth for time (‘main effect’) indicated a clear, linear decline in prevalence (EDF = 1.01, χ^2^ = 12.85, p < 0.001), the dynamics were also dependent on temperature (Figure 2B; Supplementary figure 2A for direction of change at each temperature). Compared to the main effect of time, oocyst prevalence declined at similar rates at 24 DTR 9°C (EDF = 0.07, p = 0.814; Supplementary figure 2A) and increased modestly at 28 DTR 9°C (EDF = 1.00, χ^2^ = 5.92, p = 0.015). The clearest difference from the main effect was noted at 20 DTR 9°C where prevalence increased over time before eventually leveling out at 15 dpbm (EDF = 2.02, χ^2^ = 17.02, p < 0.001) (Supplementary figure 2A).

**Figure 2.**
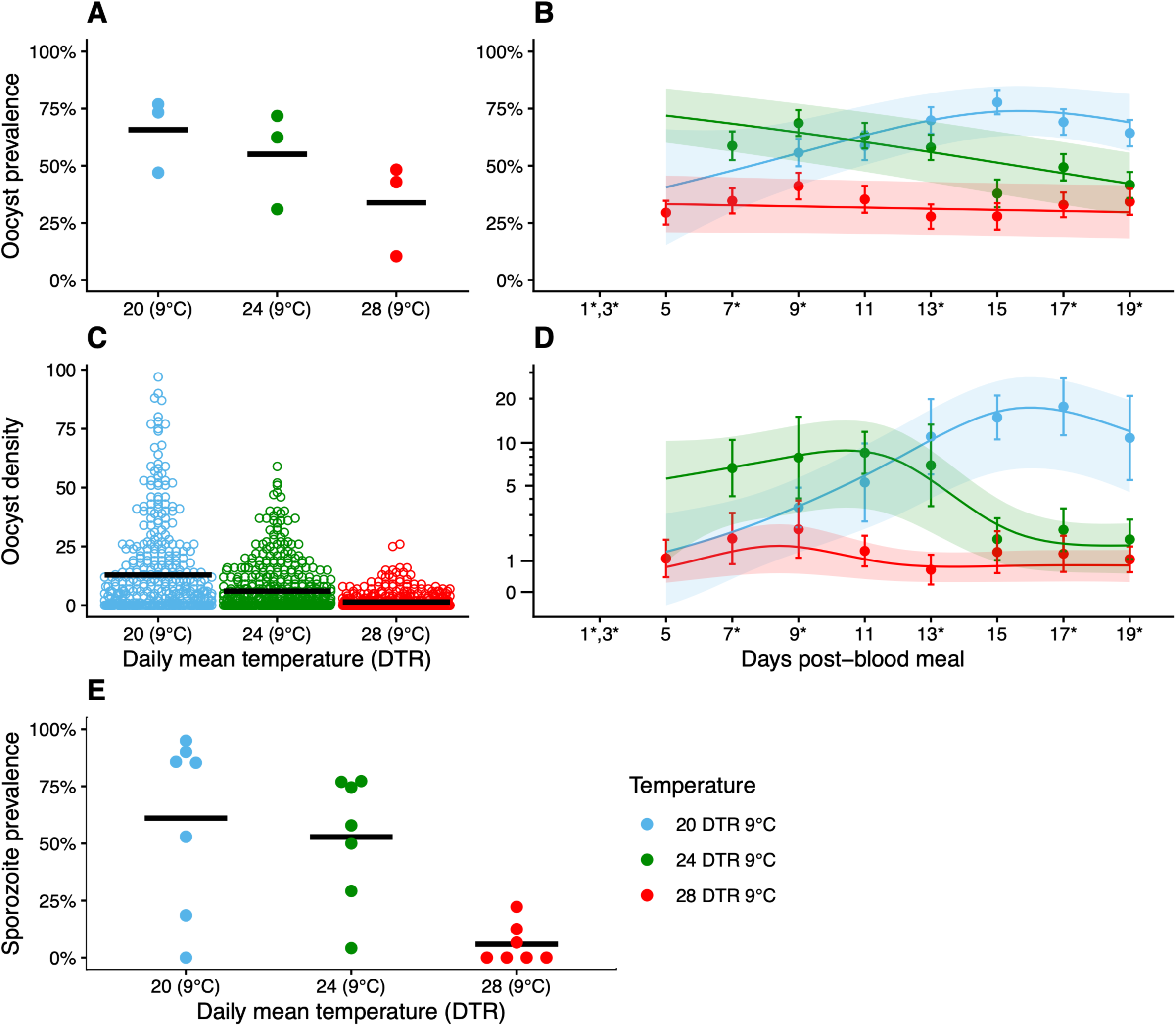
Dynamics of *P. falciparum* development in the midguts of *An. stephensi* demonstrate temperature-specific trends in oocyst prevalence (**A**), with non-linearity particularly apparent in oocyst densities (**B**). Colored lines and shaded areas indicate estimated marginal means and SE respectively. * Indicate time points where samples were collected for RNAseq: transcriptomes were characterized on 1, 3, 7, 9 and 13 dpbm for both midguts and carcasses, and additionally at 17 and 19 dpbm for carcasses only. Because oocysts are not visible at 1 and 3 dpbm at either of the three temperatures, and at 7 dpbm specifically for 20 DTR 9°C, samples were collected from mosquitoes of unknown infection status at these groups. Also see supplementary figure 1 for trends in each replicate.

Replicate effects on mean oocyst densities per midgut were again substantial (χ^2^ = 163.5, p<0.001), with lowest overall densities in replicate 1 (1.8 ± 0.5 oocysts per midgut) compared to replicates 2 (9.18 ± 2.21 oocysts per midgut) and 3 (8.19 ± 1.51 oocysts per midgut) (Supplementary figure 1B). Temperature had a strong influence on oocyst density (χ^2^ = 208, p<0.001) with higher oocyst densities observed at 20 DTR 9°C overall (vs. 24 DTR 9°C: β = −0.62, p < 0.001; vs. 28 DTR 9°C: β = −2.02, p < 0.001) (Figure 2C). While overall mean oocyst densities per midgut (‘main effect’) increased linearly over time (EDF = 1.00, χ^2^ = 24.70, p < 0.001), the temporal dynamics at the three temperatures showed clear, non-linear deviations from the main effect compared to prevalence (Figure 2D; Supplementary figure 2B). At 20 DTR 9°C, densities changed in a non-linear manner with an initial increase followed by a plateau at ∼15-17 dpbm and decline later in infection (EDF = 2.54, χ^2^ = 16.46, p = 0.008; Supplementary figure 2B). At 24 DTR 9°C, oocyst densities were initially higher than at cooler temperatures, increased modestly to 11 dpi, and then declined at later dpbm (EDF = 2.26, χ^2^ = 84.93, p < 0.001). Trends were more complex at 28 DTR 9°C with earlier decline in densities followed by partial recovery (EDF = 3.32, χ^2^ = 35.82, p < 0.001) (Supplementary figure 2B). Together, these analyses demonstrate that temperature strongly shapes the temporal dynamics of parasite establishment and growth within mosquitoes, with particularly pronounced effects on oocyst density in midguts after ∼13 dpbm.

Finally, although our initial goal of characterizing the salivary gland transcriptomes of these mosquitoes meant we were unable to determine sporozoite prevalence in these mosquitoes, we supplemented our data in the midgut (Figures 2A-D) with data from seven independent infections where sporozoite prevalence was measured in the salivary glands (Figure 2E). After adjusting for variation between replicates (χ^2^ = 11, p<0.001), infections in the salivary glands declined with temperature (χ^2^ = 40.5, p<0.001), with prevalence comparable between 20 DTR 9°C and 24 DTR 9°C (β = −0.38, p = 0.4) but significantly higher at 20 DTR 9°C than 28 DTR 9°C (β = −3.95, p < 0.001). Taken together, these results show that infections in the midguts generally translate to successful invasion of the salivary glands by sporozoites at the two cooler temperatures of 20 and 24 DTR 9°C, but not 28 DTR 9°C, where invasion increases with higher overall levels of infection in the midguts (Supplementary figure 2C), which in turn is positively associated with gametocyte density in the infectious blood meal ^22^.

### Influence of temperature, time and *P. falciparum* infection on variation in global gene expression in the midguts and carcasses of *An. stephensi*

Principle Component Analysis (PCA) plots were generated to visualize the most significant contributors to RNA-sequencing and count data variance. From these summary plots, PCA revealed that transcriptional variation was driven primarily by tissue type and time post-blood meal (Figure 3A), with early time points forming distinct clusters. In carcasses, samples clustered largely by time and temperature with minimal separation by infection status (Figure 3B), whereas midgut samples showed clearer infection-associated separation, particularly at later time points (Figure 3C).

**Figure 3.**
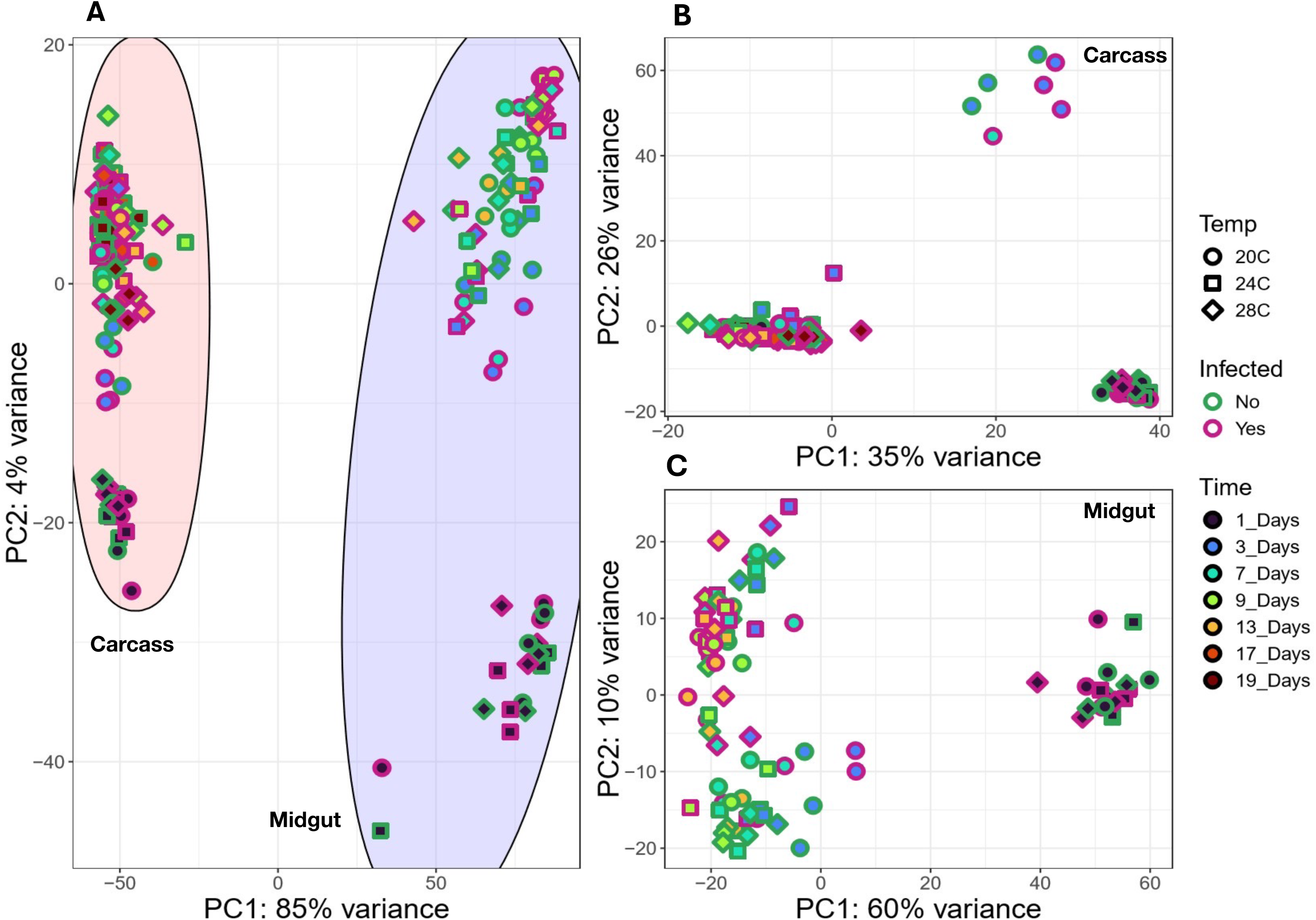
PCA plots of the influence of temperature, time and *P. falciparum* infection on variation in global gene expression of *An. stephensi* in the carcasses and midguts together (**A**), or separately for the carcasses (**B**) and midguts (**C**). Note how Day 1 time-points in both tissues cluster well together, separate from other time-points, regardless of temperature and infection status. In carcasses, the only other apparent clustering was the potentially delayed reaction at 3 dpbm the coolest temperature of 20 DTR 9°C.

The LRT models indicated that the infection parameter contributed minimally to variation in gene expression relative to temperature and time, which accounted for substantially larger variation in the transcriptome. In carcass tissues, only 18 genes showed statistically significant effects associated with infection or interactions involving infection (Figure 4). In contrast, the temperature and time experimental factors were associated with substantially larger transcriptional responses, with 1,950 and 4,188 genes identified as statistically significant, respectively. Repeating this analysis in gut tissues yielded similar patterns, with 169, 1,323, and 3,884 genes showing significant effects attributable to infection, temperature, and time, respectively. When genes associated with each factor were treated as distinct sets and their intersections visualized using UpSetR plots, the majority of genes in the infection-associated set overlapped with genes associated with temperature and/or time (Figure 4). Only one gene was identified as unique to the infection parameter in both midguts and carcasses- ASTEI02992 (a TIMELESS-domain containing protein, involved in genome stability and circadian rhythm) and ASTEI07901 (an uncharacterized protein) respectively.

**Figure 4.**
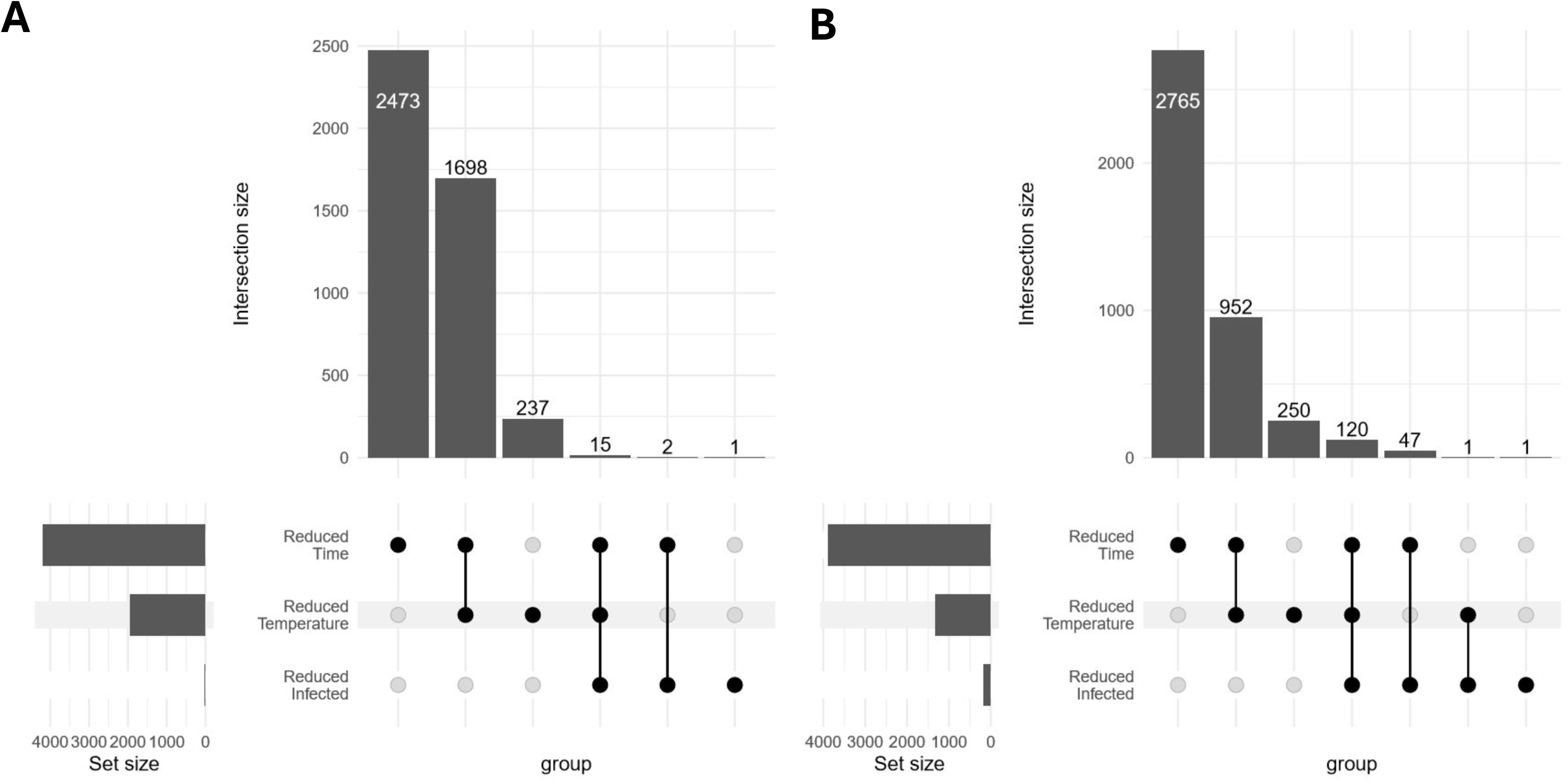
UpSetR plots of (**A)** carcasses and (**B**) midguts showing intersections of genes showing significant effects (likelihood ratio test, p<0.05) across three reduced models (Reduced Temperature, Reduced Infection, and Reduced Time; see Methods). Horizontal bars (left) indicate the total number of significant genes associated with each reduced model, while vertical bars (top) show the size of intersections between models. In (**A**), time was the dominant contributor to transcriptional variation in carcasses, with the Reduced Time model identifying 4188 genes (2473 + 1698 + 15 + 2). Of these, 2473 genes showed time-associated effects without additional contributions from temperature or infection, 1698 genes reflected combined effects of time and temperature, 2 genes showed combined effects of time and infection, and 15 genes were influenced by all three factors. Temperature was the next largest contributor (1950 genes: 1698 + 237 + 15), whereas infection contributed to a comparatively small number of genes (18). A similar pattern was observed in midguts (**B**), where time and temperature were the primary drivers of transcriptional variation. However, a larger number of genes (n = 120) showed combined effects of time, temperature, and infection in midguts compared to carcasses. For downstream analyses, we focused on genes influenced by all three factors, comprising 15 genes in carcasses and 120 genes in midguts.

### Temperature modifies the expression dynamics of genes associated with immunity, reproduction, physiology, and behavior in *P. falciparum*-infected mosquitoes

Co-expression clustering of the 15 genes identified in carcasses revealed distinct temporal expression profiles influenced by temperature (Figure 5; Table 1). Based on the primary split in the hierarchical clustering dendrogram, the smaller Clusters 1 and 2 exhibited more variable or sustained expression patterns, indicating heterogeneous transcriptional responses among infection-associated genes in carcass tissues (Figure 5). For instance, cluster 2 genes displayed sustained upregulation from 3 dpbm onward, although this response was reduced at 28 °C DTR 9°C. In both clusters, the majority of genes encode for uncharacterized proteins, aside from Aminopeptidase N (APN) in the second -largest cluster which has been suggested to be a *P. falciparum* receptor ^48^. The largest group of Cluster 3 comprised 10 genes showing increased expression at later time points, with higher temperatures generally amplifying magnitude of gene expression (Figure 5). Although these genes were primarily associated with metabolic processes (e.g. MFS transporter proteins, inositol oxygenase, F-box proteins), a subset included putative immune-related genes in Lysozyme C2 (ASTEI01308) which was recently shown to antagonize *P. falciparum* development in the midguts ^49^, and a scavenger receptor B family member “Croquemort” that reduces ookinete invasion and development of *P. berghei* in *An. gambiae* ^50^.

**Figure 5:**
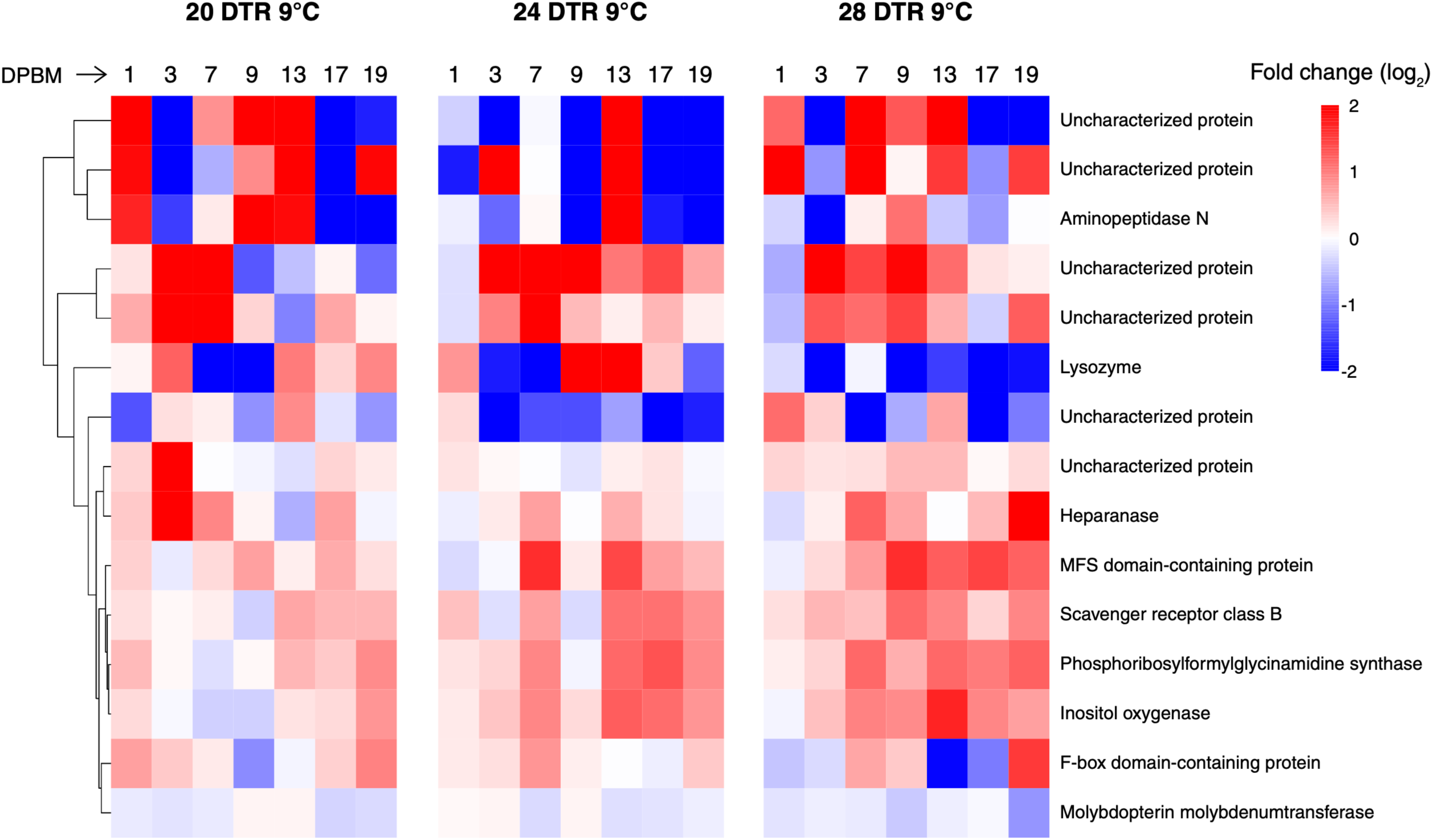
Heatmap showing expression patterns of genes in carcasses of *P. falciparum*-infected *An. stephensi* whose expression is jointly influenced by time, temperature, and infection. Color intensity represents log₂ fold-change in gene expression relative to matched controls. The dendrogram groups genes with similar expression dynamics across time, temperature, and infection, suggesting coordinated biological responses. Refer to Table 1 for corresponding VectorBase and UniProt accession numbers. Abbreviations: DPBM, days post-blood meal; MFS, Major Facilitator Superfamily.

**Table 1.**
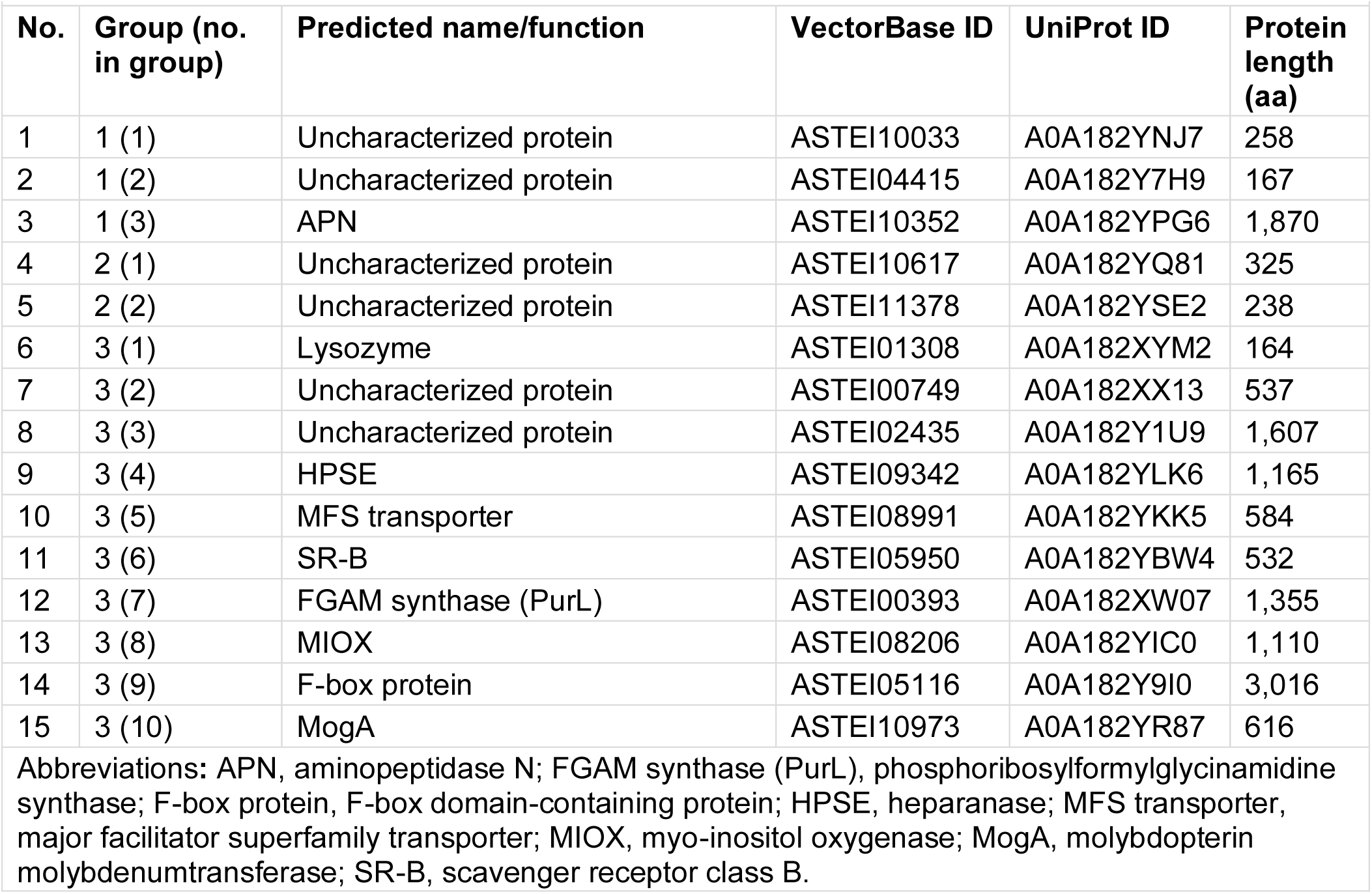
Annotations of genes differentially expressed in carcasses of *P. falciparum* infected *An. stephensi* at 1, 3, 7, 9, 13, 17 and 19 dpbm and daily mean temperatures of 20°C, 24°C and 28°C with daily fluctuations totaling 9°C (20 DTR 9°C, 24 DTR 9°C, 28 DTR 9°C). For expression dynamics, refer to Figure 1. “Group (No.)” depicts order of genes in Figure 1 by group. Abbreviations: APN, aminopeptidase N; FGAM synthase (PurL), phosphoribosylformylglycinamidine synthase; F-box protein, F-box domain-containing protein; HPSE, heparanase; MFS transporter, major facilitator superfamily transporter; MIOX, myo-inositol oxygenase; MogA, molybdopterin molybdenumtransferase; SR-B, scavenger receptor class B

| Table 1 |  |  |  |  |  |
| --- | --- | --- | --- | --- | --- |
| No. | Group (no. in group) | Predicted name/function | VectorBase ID | UniProt ID | Protein length (aa) |
| 1 | 1 (1) | Uncharacterized protein | ASTEI10033 | A0A182YNJ7 | 258 |
| 2 | 1 (2) | Uncharacterized protein | ASTEI04415 | A0A182Y7H9 | 167 |
| 3 | 1 (3) | APN | ASTEI10352 | A0A182YPG6 | 1,870 |
| 4 | 2 (1) | Uncharacterized protein | ASTEI10617 | A0A182YQ81 | 325 |
| 5 | 2 (2) | Uncharacterized protein | ASTEI11378 | A0A182YSE2 | 238 |
| 6 | 3 (1) | Lysozyme | ASTEI01308 | A0A182XYM2 | 164 |
| 7 | 3 (2) | Uncharacterized protein | ASTEI00749 | A0A182XX13 | 537 |
| 8 | 3 (3) | Uncharacterized protein | ASTEI02435 | A0A182Y1U9 | 1,607 |
| 9 | 3 (4) | HPSE | ASTEI09342 | A0A182YLK6 | 1,165 |
| 10 | 3 (5) | MFS transporter | ASTEI08991 | A0A182YKK5 | 584 |
| 11 | 3 (6) | SR-B | ASTEI05950 | A0A182YBW4 | 532 |
| 12 | 3 (7) | FGAM synthase (PurL) | ASTEI00393 | A0A182XW07 | 1,355 |
| 13 | 3 (8) | MIOX | ASTEI08206 | A0A182YIC0 | 1,110 |
| 14 | 3 (9) | F-box protein | ASTEI05116 | A0A182Y9I0 | 3,016 |
| 15 | 3 (10) | MogA | ASTEI10973 | A0A182YR87 | 616 |
| Abbreviations: APN, aminopeptidase N; FGAM synthase (PurL), phosphoribosylformylglycinamide synthase; F-box protein, F-box domain-containing protein; HPSE, heparanase; MFS transporter, major facilitator superfamily transporter; MIOX, myo-inositol oxygenase; MogA, molybdopterin molybdenumtransferase; SR-B, scavenger receptor class B. |  |  |  |  |  |

Of the 120 midgut genes whose expression was jointly influenced by *P. falciparum* infection, temperature, and time, expression was generally lower in infected mosquitoes than in controls (Supplementary figure 4A). This downregulation was strongest at 24 DTR 9°C [mean log2 fold change vs. controls = −1.30; 95% bootstrap CI: −1.47 to −1.13], followed by 28 DTR 9°C [−0.75; −0.91 to −0.59] and 20 DTR 9°C [−0.48; −0.66 to −0.29]. These temperature-specific differences reflected distinct temporal expression profiles when fold changes were averaged across the 120 genes at each temperature (Supplementary figure 4B). Except at 3 dpbm, mean expression was lower in infected mosquitoes at the two warmer temperatures, 24 and 28 DTR 9°C. By contrast, expression at 20 DTR 9°C was more heterogeneous, with higher expression at 1, 3, and 7 dpbm, followed by a decline at later time points.

Co-expression clustering of the 120 genes identified four expression clusters after the primary split in the hierarchical dendrogram (Figure 6; Supplementary figure 3C; Supplementary figure 4). The largest cluster, Cluster 1, contained 87 genes and largely explained the overall temporal trend across all 120 genes (Supplementary figure 3A,C). At 20 DTR 9°C, genes in this cluster showed higher expression in infected mosquitoes at 1 dpbm and again at 7 dpbm, with weaker infection-associated changes at 3 dpbm. At 24 and 28 DTR 9°C, expression was generally lower in infected mosquitoes, apart from transient upregulation at 3 dpbm, and declined sharply at later time points.

**Figure 6:**
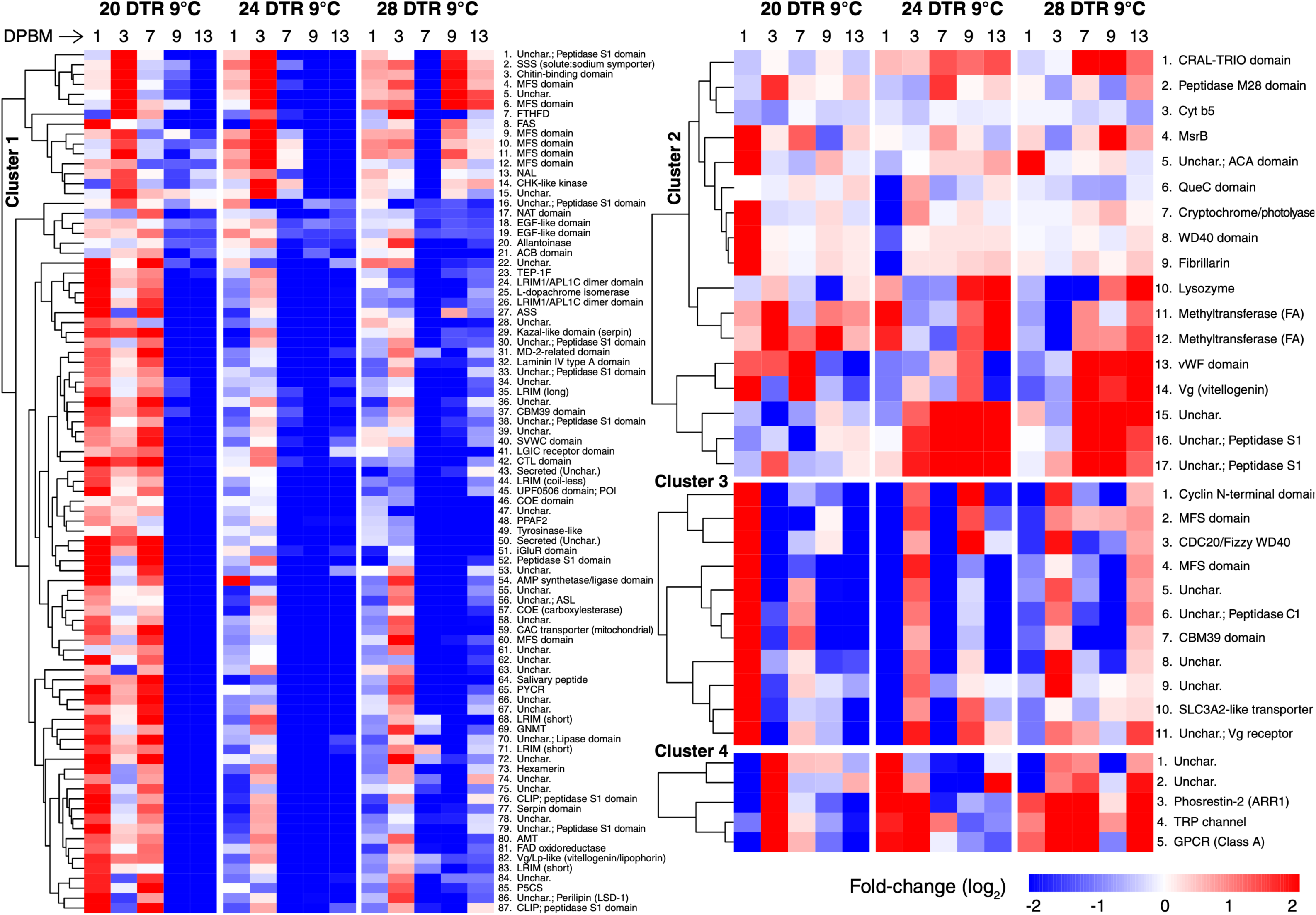
Heatmap showing expression patterns of genes in midguts of *P. falciparum*-infected *An. stephensi* whose expression is jointly influenced by time, temperature, and infection. Color intensity represents log₂ fold-change in gene expression relative to matched controls. The dendrogram groups genes with similar expression dynamics across time, temperature, and infection, possibly reflecting coordinated biological responses. For visual clarity, four expression groups were defined based on the primary split in the hierarchical clustering dendrogram; the original heatmap is shown in Supplementary Figure 2. *Vitellogenin 1 and Methyltransferase (FA-type) were excluded from downstream analysis as potential fragments of the preceding genes. Abbreviations: DPBM, days post-bloodmeal; refer to Table 2 for full list of abbreviations and corresponding VectorBase and UniProt accession numbers.

**Table 2.** Annotations of genes differentially expressed in midguts of *P. falciparum* infected *An. stephensi* at 1, 3, 7, 9 and 13 dpbm and daily mean temperatures of 20°C, 24°C and 28°C with daily fluctuations totaling 9°C (20 DTR 9°C, 24 DTR 9°C, 28 DTR 9°C). For expression dynamics, refer to Figure 2. “Group (No.)” depicts order of genes in Figure 2 by group. Abbreviations: ACA, α-carbonic anhydrase; ACB, acyl-CoA-binding domain; ALLC, allantoinase; AMT, aminomethyltransferase; AMP ligase, AMP-dependent synthetase/ligase; ARR1, arrestin-2 (phosrestin-2); ASL, argininosuccinate lyase; ASS, argininosuccinate synthase; CAC transporter, mitochondrial carnitine/acylcarnitine carrier; CBM, carbohydrate-binding module (family 39); CDC20/Fizzy, cell division cycle protein 20/Fizzy; CHK-like kinase, checkpoint kinase-like protein; CLIP domain, cysteine-rich regulatory domain; COE, carboxylesterase; CRAL-TRIO, cellular retinaldehyde-binding protein/triple function domain; CTL, C-type lectin; Cyt b5, cytochrome b5; EGF- like, epidermal growth factor-like domain; FAD oxidoreductase, flavin adenine dinucleotide- dependent oxidoreductase; FAS, fatty acid synthase; FA-type methyltransferase, fatty acid methyltransferase; Fibrillarin, nucleolar rRNA methyltransferase; FTHFD, formyltetrahydrofolate dehydrogenase; GNMT, glycine N-methyltransferase; GPCR, G protein-coupled receptor (Class A); Hexamerin, hexamerin storage protein; iGluR, ionotropic glutamate receptor; LGIC, ligand-gated ion channel; LRIM, leucine-rich repeat immune protein; MD-2-related, MD-2-related lipid-recognition domain; MFS, major facilitator superfamily transporter; MsrB, methionine sulfoxide reductase B; NAL, N-acetylneuraminate lyase-like protein; NAT, N-acetyltransferase; P5CS, Δ¹-pyrroline-5-carboxylate synthase; POI, Phenoloxidase inhibitor protein; PPAF2, prophenoloxidase-activating factor 2; PYCR, pyrroline-5-carboxylate reductase; QueC, Queuosine 5’-phosphate N-glycosylase/hydrolase; SLC3A2-like, solute carrier family 3 member 2-like transporter; Serpin, serine protease inhibitor; SSS, solute:sodium symporter; SVWC, von Willebrand factor type C-like domain; TEP-1F, thioester- containing protein 1 (full-length); TRP channel, transient receptor potential channel; Tyrosinase-like, phenoloxidase-related enzyme; UPF0506, conserved protein domain family; Vg, vitellogenin; Vg/Lp- like, vitellogenin/lipophorin-like protein; VWFD, von Willebrand factor D domain; WD40, WD40 repeat domain; Unchar., uncharacterized protein annotated based on predicted domains.

| Table 2 |  |  |  |  |  |
| --- | --- | --- | --- | --- | --- |
| No. | Group<br>(no. in<br>group) | Predicted name/function | VectorBase ID | UniProt ID | Protein<br>length<br>(aa) |
| 1 | 1 (1) | Unchar.; Peptidase S1 domain | ASTEI05101 | A0A182Y9G5 | 393 |
| 2 | 1 (2) | SSS (solute:sodium symporter) | ASTEI00312 | A0A182XVS6 | 1,214 |
| 3 | 1 (3) | Chitin-binding domain | ASTEI04105 | A0A182Y6L9 | 742 |
| 4 | 1 (4) | MFS domain | ASTEI06449 | A0A182YDB3 | 552 |
| 5 | 1 (5) | Unchar. | ASTEI04468 | A0A182Y7N2 | 217 |
| 6 | 1 (6) | MFS domain | ASTEI08991 | A0A182YKK5 | 584 |
| 7 | 1 (7) | FTHFD | ASTEI03755 | A0A182Y5L9 | 936 |
| 8 | 1 (8) | FAS | ASTEI08723 | A0A182YJT7 | 2,386 |
| 9 | 1 (9) | MFS domain | ASTEI03931 | A0A182Y645 | 952 |
| 10 | 1 (10) | MFS domain | ASTEI01702 | A0A182XZR6 | 1,349 |
| 11 | 1 (11) | MFS domain | ASTEI04280 | A0A182Y744 | 590 |
| 12 | 1 (12) | MFS domain | ASTEI05556 | A0A182YAS0 | 555 |
| 13 | 1 (13) | NAL | ASTEI01563 | A0A182XZC7 | 239 |
| 14 | 1 (14) | CHK-like kinase | ASTEI00901 | A0A182XXG5 | 809 |
| 15 | 1 (15) | Unchar. | ASTEI02758 | A0A182Y2S2 | 316 |
| 16 | 1 (16) | Unchar.; Peptidase S1 domain | ASTEI08024 | A0A182YHT8 | 270 |
| 17 | 1 (17) | NAT domain | ASTEI09896 | A0A182YN61 | 910 |
| 18 | 1 (18) | EGF-like domain | ASTEI02031 | A0A182Y0P5 | 1,290 |
| 19 | 1 (19) | EGF-like domain | ASTEI09048 | A0A182YKR2 | 132 |
| 20 | 1 (20) | Allantoinase | ASTEI01103 | A0A182XY17 | 505 |
| 21 | 1 (21) | ACB domain | ASTEI01388 | A0A182XYV2 | 111 |
| 22 | 1 (22) | Unchar. | ASTEI07844 | A0A182YHA8 | 211 |
| 23 | 1 (23) | TEP-1F | ASTEI08432 | A0A182YIZ6 | 2,611 |
| 24 | 1 (24) | LRIM1/APL1C dimerize domain | ASTEI09290 | A0A182YLF4 | 514 |
| 25 | 1 (25) | L-dopachrome isomerase | ASTEI00271 | A0A182XVN5 | 1,300 |
| 26 | 1 (26) | LRIM1/APL1C dimerize domain | ASTEI02571 | A0A182Y285 | 577 |
| 27 | 1 (27) | ASS | ASTEI03294 | A0A182Y4A8 | 424 |
| 28 | 1 (28) | Unchar. | ASTEI02560 | A0A182Y274 | 573 |
| 29 | 1 (29) | Kazal-like domain (serpin) | ASTEI01956 | A0A182Y0H0 | 63 |
| 30 | 1 (30) | Unchar.; Peptidase S1 domain | ASTEI06602 | A0A182YDR6 | 1,437 |
| No. | Group<br>(no. in<br>group) | Predicted name/function | VectorBase ID | UniProt ID | Protein<br>length<br>(aa) |
| 31 | 1 (31) | MD-2-related domain | ASTEI02361 | A0A182Y1M5 | 163 |
| 32 | 1 (32) | Laminin IV type A domain | ASTEI07094 | A0A182YF58 | 369 |
| 33 | 1 (33) | Unchar.; Peptidase S1 domain | ASTEI04157 | A0A182Y6S1 | 366 |
| 34 | 1 (34) | Unchar. | ASTEI01776 | A0A182XZZ0 | 1,028 |
| 35 | 1 (35) | LRIM (long) | ASTEI02567 | A0A182Y281 | 556 |
| 36 | 1 (36) | Unchar. | ASTEI02557 | A0A182Y271 | 83 |
| 37 | 1 (37) | CBM39 domain | ASTEI07181 | A0A182YFE5 | 561 |
| 38 | 1 (38) | Unchar.; Peptidase S1 domain | ASTEI02219 | A0A182Y183 | 1,348 |
| 39 | 1 (39) | Unchar. | ASTEI02208 | A0A182Y172 | 126 |
| 40 | 1 (40) | SVWC domain | ASTEI07751 | A0A182YH15 | 114 |
| 41 | 1 (41) | LGIC receptor domain | ASTEI05325 | A0A182YA39 | 567 |
| 42 | 1 (42) | CTL domain | ASTEI10533 | A0A182YPZ7 | 141 |
| 43 | 1 (43) | Secreted (Unchar.) | ASTEI02815 | A0A182Y2X9 | 114 |
| 44 | 1 (44) | LRIM (coil-less) | ASTEI02267 | A0A182Y1D1 | 428 |
| 45 | 1 (45) | UPF0506 domain; POI | ASTEI06661 | A0A182YDX5 | 336 |
| 46 | 1 (46) | COE domain | ASTEI00521 | A0A182XWD<br>5 | 634 |
| 47 | 1 (47) | Unchar. | ASTEI07180 | A0A182YFE4 | 251 |
| 48 | 1 (48) | PPAF2 | ASTEI08935 | A0A182YKE9 | 433 |
| 49 | 1 (49) | Tyrosinase-like | ASTEI10663 | A0A182YQC7 | 4,277 |
| 50 | 1 (50) | Secreted (Unchar.) | ASTEI06547 | A0A182YDL1 | 137 |
| 51 | 1 (51) | iGluR domain | ASTEI07031 | A0A182YEZ5 | 527 |
| 52 | 1 (52) | Peptidase S1 domain | ASTEI02562 | A0A182Y276 | 490 |
| 53 | 1 (53) | Unchar. | ASTEI09405 | A0A182YLR9 | 790 |
| 54 | 1 (54) | AMP synthetase/ligase domain | ASTEI07956 | A0A182YHM0 | 304 |
| 55 | 1 (55) | Unchar. | ASTEI01775 | A0A182XZY9 | 97 |
| 56 | 1 (56) | Unchar.; ASL | ASTEI01887 | A0A182Y0A1 | 478 |
| 57 | 1 (57) | COE (carboxylesterase) | ASTEI08772 | A0A182YJY6 | 609 |
| 58 | 1 (58) | Unchar. | ASTEI01551 | A0A182XZB5 | 188 |
| 59 | 1 (59) | CAC transporter (mitochondrial) | ASTEI07156 | A0A182YFC0 | 395 |
| 60 | 1 (60) | MFS domain | ASTEI01255 | A0A182XYG9 | 544 |
| No. | Group<br>(no. in<br>group) | Predicted name/function | VectorBase ID | UniProt ID | Protein<br>length<br>(aa) |
| 61 | 1 (61) | Unchar. | ASTEI04222 | A0A182Y6Y6 | 143 |
| 62 | 1 (62) | Unchar. | ASTEI08904 | A0A182YKB8 | 258 |
| 63 | 1 (63) | Unchar. | ASTEI04061 | A0A182Y6H5 | 134 |
| 64 | 1 (64) | Salivary peptide | ASTEI09551 | A0A182YM65 | 45 |
| 65 | 1 (65) | PYCR | ASTEI09302 | A0A182YLG6 | 233 |
| 66 | 1 (66) | Unchar. | ASTEI04108 | A0A182Y6M2 | 59 |
| 67 | 1 (67) | Unchar. | ASTEI09404 | A0A182YLR8 | 247 |
| 68 | 1 (68) | LRIM (short) | ASTEI01381 | A0A182XYU5 | 463 |
| 69 | 1 (69) | GNMT | ASTEI00591 | A0A182XWK<br>5 | 288 |
| 70 | 1 (70) | Unchar.; Lipase domain | ASTEI01076 | A0A182XXZ0 | 545 |
| 71 | 1 (71) | LRIM (short) | ASTEI01383 | A0A182XYU7 | 479 |
| 72 | 1 (72) | Unchar. | ASTEI08774 | A0A182YJY8 | 410 |
| 73 | 1 (73) | Hexamerin | ASTEI05387 | A0A182YAA1 | 1,289 |
| 74 | 1 (74) | Unchar. | ASTEI09946 | A0A182YNB0 | 471 |
| 75 | 1 (75) | Unchar. | ASTEI06949 | A0A182YER3 | 330 |
| 76 | 1 (76) | CLIP domain; peptidase S1 domain | ASTEI08921 | A0A182YKD5 | 372 |
| 77 | 1 (77) | Serpin domain | ASTEI07530 | A0A182YGE4 | 506 |
| 78 | 1 (78) | Unchar. | ASTEI07109 | A0A182YF73 | 901 |
| 79 | 1 (79) | Unchar.; Peptidase S1 domain | ASTEI08922 | A0A182YKD6 | 1,137 |
| 80 | 1 (80) | AMT | ASTEI09943 | A0A182YNA7 | 414 |
| 81 | 1 (81) | FAD oxidoreductase | ASTEI09934 | A0A182YN98 | 378 |
| 82 | 1 (82) | Vg/Lp-like (vitellogenin/lipophorin) | ASTEI00150 | A0A182XVB4 | 3,317 |
| 83 | 1 (83) | LRIM (short) | ASTEI01384 | A0A182XYU8 | 519 |
| 84 | 1 (84) | Unchar. | ASTEI09532 | A0A182YM46 | 590 |
| 85 | 1 (85) | P5CS | ASTEI11056 | A0A182YRH0 | 742 |
| 86 | 1 (86) | Unchar.; Perilipin (LSD-1) | ASTEI00476 | A0A182XW90 | 423 |
| 87 | 1 (87) | CLIP domain; peptidase S1 domain | ASTEI08920 | A0A182YKD4 | 434 |
| 88 | 2 (1) | CRAL-TRIO domain | ASTEI10314 | A0A182YPC8 | 932 |
| 89 | 2 (2) | Peptidase M28 domain | ASTEI01583 | A0A182XZE7 | 861 |
| 90 | 2 (3) | Cyt b5 | ASTEI06895 | A0A182YEK9 | 128 |
| No. | Group<br>(no. in<br>group) | Predicted name/function | VectorBase ID | UniProt ID | Protein<br>length<br>(aa) |
| 91 | 2 (4) | MsrB | ASTEI02316 | A0A182Y1I0 | 236 |
| 92 | 2 (5) | Unchar.; ACA domain | ASTEI03928 | A0A182Y642 | 914 |
| 93 | 2 (6) | QueC domain | ASTEI08521 | A0A182YJ85 | 348 |
| 94 | 2 (7) | Cryptochrome/photolyase domain | ASTEI09917 | A0A182YN81 | 542 |
| 95 | 2 (8) | WD40 domain | ASTEI05409 | A0A182YAC3 | 998 |
| 96 | 2 (9) | Fibrillarin | ASTEI06775 | A0A182YE89 | 335 |
| 97 | 2 (10) | Lysozyme | ASTEI01308 | A0A182XYM2 | 164 |
| 98 | 2 (11) | Methyltransferase (FA-type) | ASTEI09914 | A0A182YN78 | 128 |
| 99 | 2 (12) | Methyltransferase (FA-type) | ASTEI11666 | A0A182YT80 | 174 |
| 100 | 2 (13) | VWFD domain | ASTEI03705 | A0A182Y5G9 | 400 |
| 101 | 2 (14) | Vg (vitellogenin) | ASTEI11169 | A0A182YRT3 | 1,969 |
| 102 | 2 (15) | Unchar. | ASTEI05377 | A0A182YA91 | 92 |
| 103 | 2 (16) | Unchar.; Peptidase S1 domain | ASTEI08441 | A0A182YJ05 | 173 |
| 104 | 2 (17) | Unchar.; Peptidase S1 domain | ASTEI09440 | A0A182YLV4 | 163 |
| 105 | 3 (1) | Cyclin N-terminal domain | ASTEI11420 | A0A182YSI4 | 463 |
| 106 | 3 (2) | MFS domain | ASTEI02684 | A0A182Y2J8 | 721 |
| 107 | 3 (3) | CDC20/Fizzy WD40 domain | ASTEI09986 | A0A182YNF0 | 555 |
| 108 | 3 (4) | MFS domain | ASTEI00322 | A0A182XVT6 | 522 |
| 109 | 3 (5) | Unchar. | ASTEI01613 | A0A182XZH7 | 501 |
| 110 | 3 (6) | Unchar.; Peptidase C1 domain | ASTEI05380 | A0A182YA94 | 558 |
| 111 | 3 (7) | CBM39 domain | ASTEI06174 | A0A182YCI8 | 300 |
| 112 | 3 (8) | Unchar. | ASTEI01054 | A0A182XXW8 | 1,461 |
| 113 | 3 (9) | Unchar. | ASTEI00873 | A0A182XXD7 | 1,802 |
| 114 | 3 (10) | SLC3A2-like transporter | ASTEI00456 | A0A182XW70 | 148 |
| 115 | 3 (11) | Unchar.; Vg receptor | ASTEI01043 | A0A182XXV7 | 1,822 |
| 116 | 4 (1) | Unchar. | ASTEI02690 | A0A182Y2K4 | 216 |
| 117 | 4 (2) | Unchar. | ASTEI10799 | A0A182YQR3 | 213 |
| 120 | 4 (3) | Phosrestin-2 (ARR1) | ASTEI08321 | A0A182YIN5 | 383 |
| 119 | 4 (4) | TRP channel | ASTEI05371 | A0A182YA85 | 1,300 |
| 120 | 4 (5) | GPCR (Class A) | ASTEI08010 | A0A182YHS4 | 366 |
| No. | Group<br>(no. in<br>group) | Predicted name/function | VectorBase ID | UniProt ID | Protein<br>length<br>(aa) |
| Abbreviations: ACA, α-carbonic anhydrase; ACB, acyl-CoA-binding domain; ALLC, allantoinase; AMT, aminomethyltransferase; AMP ligase, AMP-dependent synthetase/ligase; ARR1, arrestin-2 (phosrestin-2); ASL, argininosuccinate lyase; ASS, argininosuccinate synthase; CAC transporter, mitochondrial carnitine/acylcarnitine carrier; CBM, carbohydrate-binding module (family 39); CDC20/Fizzy, cell division cycle protein 20/Fizzy; CHK-like kinase, checkpoint kinase-like protein; CLIP domain, cysteine-rich regulatory domain; COE, carboxylesterase; CRAL-TRIO, cellular retinaldehyde-binding protein/triple function domain; CTL, C-type lectin; Cyt b5, cytochrome b5; EGF-like, epidermal growth factor-like domain; FAD oxidoreductase, flavin adenine dinucleotide-dependent oxidoreductase; FAS, fatty acid synthase; FA-type methyltransferase, fatty acid methyltransferase; Fibrillarin, nucleolar rRNA methyltransferase; FTHFD, formyltetrahydrofolate dehydrogenase; GNMT, glycine N-methyltransferase; GPCR, G protein-coupled receptor (Class A); Hexamerin, hexamerin storage protein; iGluR, ionotropic glutamate receptor; LGIC, ligand-gated ion channel; LRIM, leucine-rich repeat immune protein; MD-2-related, MD-2-related lipid-recognition domain; MFS, major facilitator superfamily transporter; MsrB, methionine sulfoxide reductase B; NAL, N-acetylneuraminase lyase-like protein; NAT, N-acetyltransferase; P5CS, $\Delta^1$-pyrroline-5-carboxylate synthase; POI, Phenoloxidase inhibitor protein; PPAF2, prophenoloxidase-activating factor 2; PYCR, pyrroline-5-carboxylate reductase; QueC, Queuosine 5'-phosphate N-glycosylase/hydrolase; SLC3A2-like, solute carrier family 3 member 2-like transporter; Serpin, serine protease inhibitor; SSS, solute:sodium symporter; SVWC, von Willebrand factor type C-like domain; TEP-1F, thioester-containing protein 1 (full-length); TRP channel, transient receptor potential channel; Tyrosinase-like, phenoloxidase-related enzyme; UPF0506, conserved protein domain family; Vg, vitellogenin; Vg/Lp-like, vitellogenin/lipophorin-like protein; VWFD, von Willebrand factor D domain; WD40, WD40 repeat domain; Unchar., uncharacterized protein annotated based on predicted domains. |  |  |  |  |  |

Cluster 1 contained two broad functional groups. The first 14 genes were associated with metabolic functions, including MFS transport, fatty acid synthesis, proteolysis, and amino acid biosynthesis (genes 1–14; Table 2 and Figure 6). The remaining 73 genes included several genes whose *An. gambiae* orthologues are canonically associated with anti-*Plasmodium* immunity in the midgut, including components of the melanization cascade and the complement-like system ^51–54^. For example, genes associated with the LRIM1/APL1C/TEP1 complement-like system co-clustered with L-dopachrome isomerase, a key enzyme in the phenoloxidase/melanization pathway ^55,56^, suggesting coordinated expression dynamics across temperature and time (genes 23–26; Figure 6). Other genes in this cluster included CLIP-domain serine proteases, PPO-activating factor 2, prophenoloxidase, tyrosinase-like proteins ^53,54^, leucine-rich immune recognition proteins, an *An. gambiae* CTL4-like C-type lectin, and serpin regulators.

Cluster 2, the second largest cluster, contained 17 genes and showed the opposite pattern at 20 DTR 9°C compared with the two warmer temperatures. At 20 DTR 9°C, expression gradually declined over time, whereas expression increased at the same time points at 24 and especially 28 DTR 9°C (Cluster 2; Supplementary figure 3C). This cluster contained genes associated with lipid binding and transport, including vitellogenin, a putative vitellogenin-like protein with a von Willebrand factor type D domain, a CRAL-TRIO domain-containing protein, and two farnesoic acid O-methyltransferase-like genes. It also included genes associated with oxidative stress, including cytochrome b5 and methionine sulfoxide reductase B, as well as a cryptochrome/photolyase alpha/beta domain-containing protein similar to cryptochrome 1, a photoreceptor involved in circadian regulation in mosquitoes ^57^.

Cluster 3 contained 11 genes and displayed more complex, nonlinear expression dynamics, with infection-associated upregulation or downregulation depending on temperature and time (Cluster 3; Supplementary figure 3C). Excluding uncharacterized proteins, this cluster included a putative vitellogenin receptor, nutrient transport-associated genes, including a CBM39-domain protein and an SLC3A2-like transporter, and cell-cycle-associated genes, including a cyclin N-terminal domain- containing protein and a CDC20/Fizzy WD40 domain-containing protein (Cluster 3, genes 1–11; Table 2 and Figure 6).

The smallest cluster, Cluster 4, contained five genes and showed higher overall expression at 28 DTR 9°C, with broadly similar but weaker dynamics at 20 DTR 9°C (Cluster 4; Supplementary figure 3C). At both temperatures, expression peaked at 3 dpbm. However, at 28 DTR 9°C, expression declined more slowly until 9 dpbm before increasing again at 13 dpbm. At 24 DTR 9°C, expression was highest at 1 dpbm and then declined gradually across the remaining time points. Apart from two uncharacterized genes, this cluster contained genes encoding a transient receptor potential channel, a G protein-coupled receptor, and Phosrestin-2 (Cluster 4, genes 1–5; Table 2 and Figure 6). TRP channels regulate insect responses to environmental stimuli, including temperature ^58^, whereas the GPCR and Phosrestin-2 proteins are associated with photoreception ^59^.

### Temperature elicits canonical responses to *P. falciparum* infections in the midguts of *An. stephensi*

Enrichment analysis revealed no significant functional groupings among the 15 carcass genes whose expression was jointly influenced by temperature, time and *P. falciparum* infection. In contrast, the 120 midgut genes elicited by the three treatments showed strong enrichment for extracellular and proteolytic functions, consistent with roles in digestion and immune defense (Table 3). Several enriched categories corresponded to canonical anti-*Plasmodium* immune pathways, including the Toll and Imd signaling pathways and associated pattern recognition (β-1,3-glucan-binding proteins) and effector systems (LRR proteins, CLIP-domain proteases) ^60,61^ (see Supplementary datasheet 1 for corresponding *An. gambiae* orthologs and list of genes in each category). In addition to immune-related processes, enrichment highlighted genes involved in nutrient transport and core metabolic pathways, including amino acid metabolism that feed into the folate-mediated one-carbon metabolism required for nucleotide biosynthesis; the importance of folate pathways for *P. falciparum* replication is underscored by the long- standing use of antifolate drugs such as sulfadoxine–pyrimethamine ^62,63^.

**Table 3:** Enrichment analysis of 120 genes differentially expressed in the midguts of *P. falciparum*- infected *An. stephensi* midguts relative to controls. To main clarity and brevity, the above categories are reported after pooling redundant terms, using STRING database software default of “Redundancy cutoff = 0.5” (range 0-1) ^105^.

| **Table 3** |  |  |  |  |
| --- | --- | --- | --- | --- |
| **Category** | **Description** | **# genes** | **# background genes** | **FDR value** |
| UniProt Keywords | Signal | 42 | 1270 | 7.45E-10 |
| GO Cellular Component | Extracellular region | 32 | 842 | 8.6E-08 |
| SMART Domains | Trypsin-like serine protease | 14 | 167 | 0.00000196 |
| SMART Domains | Clip or disulphide knot domain | 4 | 13 | 0.0058 |
| InterPro Domains | MFS transporter superfamily | 9 | 132 | 0.0074 |
| KEGG Pathways | Toll and Imd signaling pathway | 8 | 125 | 0.0096 |
| InterPro Domains | Leucine-rich repeat, typical subtype | 8 | 111 | 0.0108 |
| SMART Domains | von Willebrand factor (vWF) type D domain | 3 | 6 | 0.0146 |
| UniProt Keywords | Disulfide bond | 14 | 416 | 0.015 |
| InterPro Domains | β-1,3-glucan-binding protein, N-terminal | 3 | 5 | 0.0181 |
| InterPro Domains | Farnesoic acid O-methyl transferase | 3 | 7 | 0.03 |
| STRING Clusters | α-amino acid metabolic process, and One carbon pool by folate | 7 | 86 | 0.0331 |

Together, these results suggest that the midgut response to infection reflects coordinated regulation of immune and metabolic pathways in a temperature- and time-dependent manner. For example, similar to how expression of TEP-1F, CTL4-like, and LRIM1/APL1C-like genes declined in infected mosquitoes as temperature increased (Figure 6; Supplementary datasheet 2), at 28 DTR 9°C, this temporal downregulation was also evident for four of eight genes assigned to the “Toll and Imd signaling pathway” category: a peptidase S1 domain-containing protein (ASTEI02562), a serpin-domain protein orthologous to *An. gambiae* SRPN11 (ASTEI07530), an uncharacterized gene (ASTEI06949), and a putative CLIP-domain serine protease orthologous to *An. gambiae* CLIPE4 (ASTEI07109) (Table 3; Figure 6; Supplementary datasheet 2). By contrast, the remaining four genes in this pathway—CLIP- domain serine proteases ASTEI04157 (*An. gambiae* CLIPA9), ASTEI06602, ASTEI08921 (*An. gambiae* CLIPB1), and ASTEI08920 (*An. gambiae* CLIPB6)—were upregulated at least transiently at 3 dpbm. Similar early upregulation was observed for other LRIM genes in the “Leucine-rich repeat” category, including ASTEI01381, ASTEI09946, and ASTEI02560.

In contrast to the immune-associated genes that tended to decline at 28 DTR 9°C, seven genes assigned to the “α-amino acid metabolic process and one-carbon pool by folate” category showed distinct temperature-dependent expression profiles (Table 3; Figure 6; Supplementary datasheet 2). These genes were generally downregulated at 24 DTR 9°C, whereas expression was higher at 20 and 28 DTR 9°C from 1 to 7 dpbm before declining at 9 and 13 dpbm. This group included argininosuccinate synthase (ASS, ASTEI03294), a likely argininosuccinate lyase (ASL, ASTEI01887), pyrroline-5-carboxylate reductase (PYCR, ASTEI09302), glycine N-methyltransferase (GNMT, ASTEI00591), aminomethyltransferase (AMT, ASTEI09943), an FAD-dependent oxidoreductase domain-containing protein (ASTEI09934), and Δ^1^-pyrroline-5-carboxylate synthase (P5CS, ASTEI11056). These genes were assigned to midgut Cluster 1, consistent with their shared expression dynamics.

The three genes containing “von Willebrand factor (vWF) type D domains” showed more heterogeneous clustering and expression patterns (Table 3; Figure 6; Supplementary datasheet 2). The *An. stephensi* vitellogenin/lipophorin-like gene (Vg/Lp-like, ASTEI00150) was assigned to Cluster 1 and showed higher expression from 1 to 7 dpbm, followed by downregulation at later time points (Supplementary figure 3C). The other two vWF domain-containing genes—a putative vitellogenin-like gene (ASTEI03705) and vitellogenin (Vg, ASTEI11169)—were assigned to Cluster 2 and showed distinct temperature-dependent trajectories (Supplementary figure 3C). At 20 DTR 9°C, both genes were upregulated from 1 to 7 dpbm before declining at later time points. At 24 DTR 9°C, expression was generally lower, except for transient upregulation at 9 dpbm. At 28 DTR 9°C, however, both genes showed sustained upregulation relative to uninfected controls at 7, 9, and 13 dpbm.

### Temperature perturbs key biological networks in *P. falciparum*-infected midguts

Network analysis of the 120 midgut genes identified a significantly interconnected STRING network of 78 proteins with more (117) interactions than expected by chance (expected = 69, p = 2.18e-07), suggesting that the infection-responsive transcriptome is not isolated but instead reflects coordinated regulation across inter-connected biological processes (Figure 7; Supplementary datasheet 3). Whereas the gene- set enrichment analysis above (Table 3) identified the dominant functional categories among the 120 genes, the network approach revealed associations among these categories that were not captured by enrichment analysis alone (Figure 7; Supplementary datasheet 1 lists genes assigned by one or both approaches). In particular, the network placed immune, amino acid/folate metabolic, lipid-transport, vitellogenin-like, sensory, and cell-cycle-associated proteins within the same connected system, suggesting that temperature-dependent infection responses may be organized around shared physiological bottlenecks rather than independent pathways.

**Figure 7:**
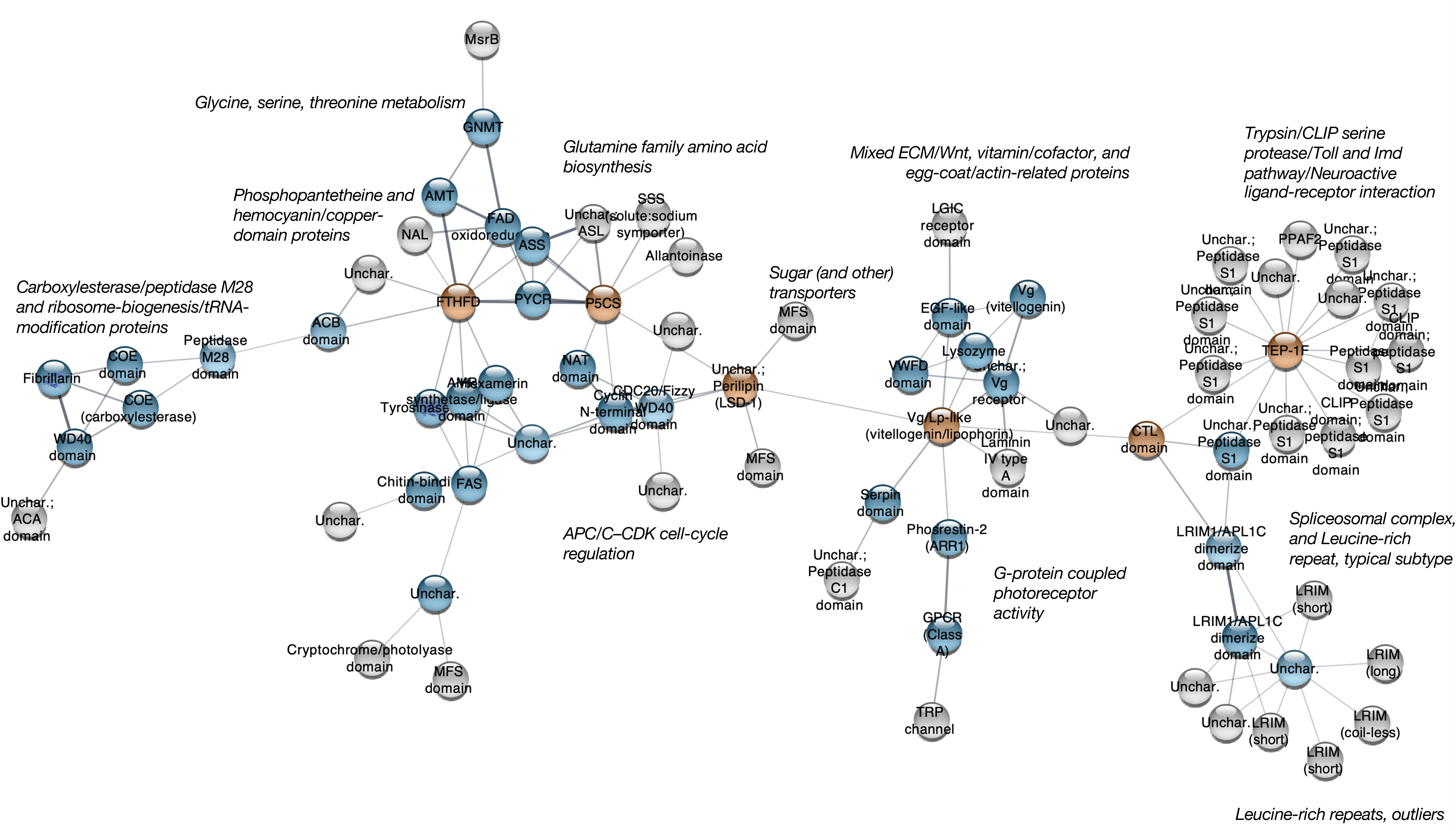
Relative influence of proteins within the STRING network of 77 differentially expressed genes in *P. falciparum*-infected *An. stephensi* midguts relative to controls. Grey edges (lines) represent predicted protein–protein associations derived from the STRING database, with edge thickness depicting interaction confidence score (range 0.4–1.0). Node color reflects ‘BetweennessCentrality’, a measure of the relative importance of proteins in maintaining network connectivity: vermilion-filled nodes indicate highly central “bottleneck” proteins, blue nodes indicate intermediate centrality, and light gray nodes indicate low centrality. Labels in italics denote the most significantly enriched (FDR<0.05) functional categories within each MCL-defined cluster (lowest FDR).

MCL clustering of the STRING network further separated the 78 interacting proteins into functional modules (Figure 7) that extended beyond the broader enrichment terms in Table 3 (Supplementary table 1). While group-wise enrichment analysis of these modules identified terms associated with immune, metabolic, reproductive, sensory, and regulatory processes, including trypsin/CLIP serine proteases, Toll and Imd signaling, leucine-rich repeat proteins, and amino acid metabolism, the network also recovered smaller modules associated with Phosphopantetheine ^64^/hemocyanin-copper domains, APC/C–CDK cell- cycle regulation, glycine/serine/threonine metabolism and photoreceptor/GPCR-related proteins (Figure 7; Supplementary table 1; Supplementary datasheet 3).

Within this network, several proteins showed high betweenness centrality (BC), identifying them as potential bottlenecks connecting otherwise distinct functional modules (Supplementary figure 6). Using a data-driven threshold of BC > mean + 1 SD among nodes with nonzero BC values (BC > 0.265), six proteins were classified as central nodes linking immune, metabolic, and lipid-transport pathways ^47^ (vermillion nodes, Figure 7). These bottleneck proteins span immune, metabolic, and reproductive functions, suggesting that infection-associated responses in the midgut may involve a shared physiological network linking parasite killing, nutrient allocation, and life-history processes. The six proteins with BC estimates >0.265 (Supplementary figure 6) were assigned to the largest Cluster 1 of the midguts (Figure 6). Consistent with the broader Cluster 1 expression profile (Supplementary figure 3C), these genes showed higher expression at 1, 3 and 7 dpbm at 20 DTR 9°C, and 3 dpbm at the two warmer temperatures, followed by reduced expression at the later time points of 9 and 13 dpbm at all three temperatures (Figure 6; Supplementary figure 7). Notable exceptions to these patterns were two proteins integral to the anti-*Plasmodium* immune response in the C-type lectin domain-containing protein, a pattern recognition receptor and ortholog of *An. gambiae* CTL-4 ^65^, and TEP-1F ^66,61^, with reduced expression at 28 DTR 9°C. Similar reduction in expression was also observed at 24 DTR 9°C for P5CS (Δ¹-pyrroline-5-carboxylate synthase), a key regulatory enzyme in the biosynthesis of proline from glutamate ^67^ and may in turn provide an energy substrate for flight ^68^. Taken together, these results suggest a clear influence of temperature and time on protein-protein interactions critical to *Plasmodium– Anopheles* association.

## Discussion

The trends in oocyst prevalence and densities clearly demonstrate the effect of temperature on parasite development rates (Figure 2). Differential gene expression identified the majority of mosquito transcriptional responses were shaped by tissue, time post-infection, and temperature (Figures 3, 4), with *P. falciparum* exerting little influence in the carcasses compared to the midguts (Figure 4). Nonetheless, our analysis identified key genes associated with vector immunity, reproduction, nutrient acquisition, metabolism and behavior (Figures 6, 7; Table 3; Supplementary table 1). Together, our results also suggest that any inference regarding the consequences of the association cannot be taken in isolation from variation in environmental variables, like temperature.

The observed dynamics of oocyst prevalence and densities generally corroborate previous studies demonstrating the effect of temperature on the dynamics of sporozoite appearance in the salivary glands ^69,70,20,21^. Since both oocyst measures are dependent on being large enough to detect microscopically, the apparent changes over time may have been due to the changes in visibility as size of oocysts increase initially due to sporozoite proliferation ^71,72^, with the subsequent decline possibly reflecting the difficulty in detecting ruptured and/or repaired midguts as sporozoites migrate to the salivary glands ^73,74^. While we initially posited that any temperature-associated mortality of infected mosquitoes during the sampling window would be expected to appear as a corresponding decline in infection prevalence especially at the two warmer temperatures, our model suggested declines were more apparent after 13 dpbm^10^. Additionally, oocyst prevalence and density displayed non-linear temperature- dependent dynamics. After the slope of oocyst densities declined at approximately 9 and 13 dpbm under 28°C and 24 DTR 9°C conditions, respectively, it increased again, particularly at the warmer temperature. However, mean oocyst densities remained lower than those observed during the preceding period. While it is possible these trends may be more apparent at 20 DTR 9°C after 19 dpbm, whether this increase is due to a sub-population of asynchronously developing oocysts ^72,75^, this may warrant further examination ^76–78^ if asynchronicity in oocyst development cascades to the rates of sporozoite migration and densities.

Relatively few genes were affected by *P. falciparum* infection in the carcasses of the mosquitoes. While this may appear inconsistent with the extensive literature describing the pathways involved in mosquito immune responses to *Plasmodium* infection ^79–81^, this could be attributed to various factors such as lower signal-noise ratio due to the sequencing depth and read length, masking by transcriptomes of non-target tissues ^82^, or our decision to use a more conservative, LRT-based framework to identify genes whose expression is consistently influenced by infection across all experimental conditions (refer to “Differential expression analysis” section for more detail). The primary source of variation in gene expression was the blood meal and the delayed transcriptional responses at 3 dpbm under the 20 DTR 9°C regime. For the remaining five sampling points until 19 dpbm, the overlapping infected and control data points suggest overall systemic transcriptional profiles were broadly similar between infected and control mosquitoes. As such, the smaller number of infection-associated genes detected here likely reflects a more robust estimate of genes consistently influenced by infection across the full experimental schedule. Taken together however, this result suggests minor influence of *P. falciparum* infections at systemic levels.

Amongst the 15 DEGs identified in the carcasses, at least three, Lysozyme C2 ^49^, Scavenger receptor B (“Croquemort” ^50^) and Aminopeptidase N ^48^ have been shown to reduce *Plasmodium* infections. However, it is unclear if their activity is dependent on temperature. For instance, dsRNA- mediated silencing of Lysozyme C2 expression led to enhanced infections of *An. stephensi* midguts with *P. falciparum* ^49^. However, our results suggest expression of the same gene, while downregulated in the carcasses at 28 DTR 9°C compared to the two cooler temperatures, showed an inverse pattern in the midguts with lower expression at 20 DTR 9°C compared to 24 and 28 DTR 9°C. As such, the expression patterns in the carcasses are more compatible with Lysozyme’s status as a cold-adapted protein^83^, while patterns in the midgut suggest an association with reduced midgut infections at the two warmer temperatures. Taken together, our results suggest tissue-specific functions of this enzyme ^82^ that is regulated by a critical environmental variable, temperature. As we and others have shown previously ^21,22,70^, sporozoite prevalence in the salivary glands is significantly reduced over the thermal gradient. Based on the expression levels in the carcasses, there appear to be few genes that are differentially expressed that explain the dynamics of sporozoite migration and sporozoite prevalence at each temperature. However, scavenger receptor B could be a suitable starting point to assess if upregulation of this gene at the two warmer temperatures and later time points reduces sporozoite invasion, similar to its role in preventing ookinete invasion earlier in the infection ^50^.

In the midguts, temperature and *P. falciparum* infection elicited expression of genes that we manually grouped into four clusters based on the structure of the dendrogram. The first and largest cluster comprised multiple genes related to canonical immune responses being activated at the site of infection in the midguts, with immune recognition proteins and pathways identified. Enrichment analysis suggested several of the 120 genes were known or predicted to be involved in mosquito immunity^61,84^. In addition, the trends in expression were positively correlated with the early stages of parasite development, suggesting their roles are likely limited to preventing invasion of the midgut epithelia by the ookinete ^61^, with the downregulation at the later stages likely due to the oocysts ability to evade immunity by binding mosquito proteins such as laminin, lysozyme c-1 and matrix metalloprotease 1 ^61,84^ (note that the list of genes also included a laminin IV type A domain-containing protein in Cluster 1^85^ and a lysozyme in Cluster 2). While downregulation of immune response genes at the later time points are consistent the aging-associated reduction in immune function ^86–88^, tight regulation of these genes may be necessary to minimize fitness costs due to toxicity resulting from activation of these effectors ^89,90^ and divert resources towards other life history processes ^91^.

For this same cluster (Cluster 1), in addition to delayed expression until 7 dpbm at 20 DTR 9°C, overall expression levels in infected mosquitoes, relative to the controls, were generally higher at 20 DTR 9°C compared to the two warmer temperatures. This was likely not due to lower expression levels of these genes in the control mosquitoes at 20 DTR 9°C, as our previous report^27^ demonstrated higher levels and diversity of gene expression in the midguts of this reference group at 20 DTR 9°C. Slower rates of mRNA turnover at the cooler temperatures ^92^ may result in transcripts persisting for a longer duration compared to the warmer temperatures, and thus, more closely spaced sampling points could reveal similar expression levels across the three temperatures after adjusting for thermal shifts in temporal dynamics; however, the transient downregulation of this sub-cluster at 3 dpbm at 20 DTR 9°C suggests this may be unlikely.

We consider two possible explanations for this finding, which, collectively, highlight the benefits of a transcriptomic approach to resolve the mechanisms mediating trade-offs between vector traits, which in turn impose selection pressure on the malaria parasites and their evolutionary responses ^93^. First, if temperature has direct effects on parasite survival, with parasite mortality being higher at warmer temperatures, the higher immune activation status at 20 DTR 9°C could reflect higher parasite exposure rates relative to the two higher temperatures ^16–19^. Examination of these responses to a range of parasite densities or genotypes from different environments may be insightful. Second, this finding may reflect life history trade-offs shaped by temperature. As temperatures warm, mosquito lifespans are shorter. Thus, infected mosquitoes may allocate resources to reproduction and survival at the warmer temperature, at the cost of immunity, to maximize fitness ^89,91^. These trade-offs may be evident in how temperature modulates the levels and temporal expression profiles of genes in the midgut clusters – while expression of the immune-related genes in cluster 1 declined with temperature and time, general opposing trends were noted for the genes in clusters 2 – 4 (and gene numbers 1-14 in cluster 1) with roles in nutrient transport and biosynthesis, oxidative stress, circadian regulation and photoreception (Figure 6).

One example of this shift is the expression of three genes– 1) the Vg/Lp ortholog in cluster 1, the yolk precursor protein Vitellogenin (Vg) in cluster 2 (and another putative Vg in VWFD domain-containing protein), and its corresponding receptor (Vg receptor) in cluster 3. While the transient upregulation of all three genes in infected mosquitoes between 1 to 7 dpbm is consistent with established roles in vitellogenesis and immunity, elevated expression of Vg and VgR at the warmer temperatures, along with the other nutrient cycling and biosynthetic genes, may reflect their broader functions in nutrient allocation, stress tolerance, lifespan regulation and fecundity (lifetime fitness) ^91,94^. Notably, dsRNA-mediated knockdown of Vg and VgR reduced survival after thermal stress in the melon fruit fly, *Zeugodacus cucurbitae* (Coquillett) ^95,96^ and the honey bee *Apis mellifera* ^97^. Although comparable studies in mosquitoes are lacking, higher expression of these genes in *P. falciparum*-infected midguts may provide a mechanistic basis to explain how infected mosquitoes show higher survival and increased resistance to environmental stressors relative to uninfected mosquitoes ^98–100^. Additionally, upregulation of the GPCR photoreceptor gene family could further benefit parasite fitness by enhancing host-seeking activity^101,102^.

Despite recent updates, the sparser annotation of the *An. stephensi* genome presents a greater hurdle to downstream analysis and interpretation than *An. gambiae* ^103,104^. Nonetheless, enrichment/overrepresentation (Table 3) and network analysis (Figure 7; Supplementary table 1) together accounted for most (104) of the 120 midgut genes (∼87% coverage). Of these 104 genes, 53 genes were mapped by both analyses, with another 26 and 25 genes unique to the enrichment and network analysis, respectively (53 + 26 + 25 = 104, Supplementary datasheet 1). Network analysis suggested influence of temperature on protein networks mediating immune, metabolic, reproductive and behavioral processes in the midguts of *P. falciparum*-infected *An. stephensi* and predicted protein ‘bottlenecks’ with disproportionate influence in bridging the various processes in a network. Although we focused on the six proteins that passed the distribution threshold, most of the proteins with non-zero estimates of BC (e.g., LRIM1/APL1C-like proteins, vitellogenin, lysozyme, serpins, and fatty acid synthase) are well- documented components of mosquito immune, metabolic, and reproductive responses to *Plasmodium* infection ^61^. Because these bottlenecks are identified from known and predicted protein–protein interactions across diverse biological contexts ^105^, they should be the first line of enquiry in assessing the mechanistic basis for how temperature influences *Plasmodium* fitness.

The high BC estimates of Lp, CTL-4 and TEP-1 are consistent with their role as regulatory bottlenecks for *Plasmodium* fitness in the midguts ^61,106^. There is substantial, indirect evidence suggesting temperature-dependent contributions of Lp and CTL4 but especially TEP-1 in regulating the fitness of *P. falciparum* and the rodent malaria species *P. berghei*: *P. falciparum*-infected mosquitoes are typically housed at ∼27°C, while *P. berghei* infections require a much cooler temperature of 20°C (DTR = 0°C) ^65^. The technical challenges associated with in vitro cultures of *P. falciparum* has meant much of our understanding of the mosquito immune repertoire is derived from the rodent parasite. In general, the rodent parasite elicits much stronger immune responses than the human parasite, with TEP-1 mediated melanization and phagocytosis central to the effector response ^106^. However, our results suggest expression levels of TEP-1, like many of the other immune genes in Cluster 1 of the midgut (Figure 6), are higher at the coolest temperature of 20 DTR 9°C, with declining levels at the intermediate temperature followed by almost complete downregulation at the warmest temperature of 28 DTR 9°C. While also true for CTL-4 ^65^ and the two LRIM1/APL1C-like proteins, the gene expression patterns are correlated with functional activity based on our previous observation of higher melanization and phagocytosis at the cooler temperatures ^107^. Considered together, any inferences of immune efficacy in the vector should consider the possibility that an effect, or the lack thereof, may be confounded by the temperature experienced by the mosquito.

However, roles for LSD-1, P5CS and FHD are not clear and warrant further examination. LSD-1 functions upstream of Lp in the Adipokinetic Hormone (AKH) signaling pathway that promotes lipid mobilization through LSD-1–mediated regulation of lipid droplet lipolysis, with lipophorin subsequently shuttling mobilized diacylglycerol through the hemolymph to support systemic energy demands ^108^. LSD-1 is also implicated in vector-pathogen systems ^109–111^. P5CS is a key regulatory enzyme in the biosynthesis of proline from glutamate (along with PYCR, gene number 65, Cluster 1, Figure 6), while FHD carries out the NADP^+^-dependent conversion of 10-formyl-tetrahydrofolate (THF) to THF necessary for purine synthesis. Disruption of a proline transport gene in *P. falciparum* (PfApiAT2) resulted in higher number of oocysts in *An. gambiae*, although they were unable to continue their development, measured as oocyst diameter, or initiate sporozoite production (sporulation) ^112^. Of the two folate transporter genes (*ft1* and *ft2*) expressed by *Plasmodium*, disrupting expression of FT2 in *P. berghei* did not affect oocyst burdens or growth rates (i.e., size) in the midguts but impaired sporulation; this defect was specific to the later stages of sporogony as parasitemia and gametocytogenesis in mice was similar to wild type parasites ^113^.

In addition to some of the limitations discussed above in reconciling the effect of temperature on vector survival and infection burdens, the relatively few infection-associated genes in the carcasses and the annotation status of the *An. stephensi* transcriptome, there are three additional caveats worth noting. First, while the transcriptomes at 1 and 3 dpbm are from mosquitoes with unknown infection status, increasing temperatures are associated with a strong decline in infection levels^22^. Therefore, our inference of gene expression patterns at these early time points may be impacted by the proportion of infected mosquitoes, particularly at 28 DTR 9°C. For instance, while TEP-1, CTL-4 and the LRIM1/APL1C-like proteins were downregulated at all three temperatures at the later time points, at 28 DTR 9°C, expression is downregulated at all the time points. While this could be due to lower infection levels at this temperature, several other genes of the immune repertoire suggested evidence of upregulation at early time points, such as four of the eight genes in the “Toll and Imd pathways” and three genes in “Leucine rich repeat” categories. These results suggest there may be selective downregulation of specific components of vector immunity. Second, in the absence of corresponding information on protein expression, our results should be taken with a degree of caution, particularly for the genes that are part of the immune repertoire and undergo further post-translational processing before they are activated. However, in general, the temperature-dependent effects on gene expression of the immune repertoire is supported by our previous observation ^107^ and those of other studies that show higher melanization and phagocytosis at cooler temperatures ^88,114,115^. Third, although our analysis of the midgut genes suggests an alteration of critical networks, the 15 genes in the carcass and lack of enrichment or networks suggest this may not extend to the systemic level. However, this is likely due to our experimental design and choice of statistical analysis resulting in a lower signal-noise ratio, as the midgut is the primary site of pathogen invasion, nutrient digestion, absorption, and is a key source of endocrine and physiological signaling that could result in systemic responses mediating immune-physiology, digestion, reproduction, and behavior. Thus, a possible solution for future studies would be to consider adjusting the sequencing depth and read length to the type of tissue and expected size of transcriptome ^82^.

## CONCLUSIONS

Temperature is a major regulator of *Plasmodium*–*Anopheles* interactions, influencing parasite development rates, and host transcriptional responses across tissues and time. Although temperature and time were the primary contributors to variation in the transcriptome, infection elicited modest but canonical anti-*Plasmodium* responses in the midgut. Rather than inducing widespread changes in gene expression, our results suggest *P. falciparum* infections influence coordinated networks linking reproduction, immunity, metabolism, and behaviour, mediated in part by a small number of highly connected “bottleneck” proteins. We suggest these proteins should be key starting points to identify novel mechanisms that prevent (or promote) *Plasmodium* fitness in the context of a key ecological determinant in temperature but also re-evaluate the extent to which cross-species comparisons are confounded by experimental temperature regimes. Temperature influenced not only the magnitude but also the organization of mosquito responses to infection, with potential consequences for resource allocation and life-history trade-offs. Midgut responses suggest two interacting mechanisms: temperature-driven differences in parasite survival shaping immune activation, and life-history trade- offs that prioritize reproduction and survival over immunity at warmer temperatures. More broadly, our study highlights the value of integrating temporal and environmental dimensions into transcriptomic to resolve the mechanistic basis of vector competence and its sensitivity to changing climates.

## Supplementary datasheet legends

**Supplementary datasheet 1**: Sortable list of genes whose expression was jointly influenced by time, temperature, and infection in the midguts of *P. falciparum*-infected *An. stephensi*. Columns A-G list the genes identified in this study and stated in Table 2 of the main manuscript. Orthologues of these genes from *An. gambiae* (PEST) are listed in Columns H-L, with column names starting with “AGAM…” listing corresponding VectorBase and UniProt accession numbers and genes names where available; the column titled “similarity (bitscore)” indicating the strength of sequence alignment between proteins from the two species, with higher values indicating greater sequence similarity. Columns M and N respectively state which *An. stephensi* gene was included in the enrichment analysis (Table 3) and network (Figure 7); columns titled “in.STRING.enrichment?” and (in.STRING.network?) lists genes included (TRUE/FALSE), with values of TRUE in both columns depicting overlap in both analyses. Finally, columns P-R lists the labels/names, along with the key metrics used to annotate the network (Figure 7): while “node.degree” reflects the number of direct interaction between proteins, with higher values suggested highly connected ‘hub’ proteins, “BetweennessCentrality”, by contrast, identifies proteins that act as bridges within the network linking distinct functional groups and potentially mediating crosstalk between biological processes.

**Supplementary datasheet 2**: This datasheet extends Table 3 by including the genes in each enrichment category.

**Supplementary datasheet 3**: Sortable datasheets listing performance and graph attributes and other metadata that were used to build and annotate the network in Figure 7 via the stringApp (v2.2.0) in Cytoscape (v3.10.4). The first datasheet titled “Edge (PPI) table” describes the metrics used to identify predicted or known PPI between the genes in our dataset, with confidence score (0–1) indicating the strength of supporting evidence for each PPI (derived from integrated sources such as experimental data, co-expression, and orthology); higher values reflecting greater confidence, as indicated by the thickness of grey lines in Figure 7. Note that these scores represent the likelihood of an association and should not be interpreted as a measure of the strength or affinity of a physical interaction. The second datasheet (“Node attributes”) lists the functional and topological information for each protein, and their position in the respective MCL clusters assigned by STRING (Column B, “ mclCluster”), as well as the “BetweennessCentrality” values (Column D) discussed earlier assigned as node color in the network (Figure 7).

**Supplementary datasheet 4**: This datasheet extends Supplementary table 1 by listing the genes enriched in each MCL cluster.

## Supplementary figures

**Supplementary figure 1:**
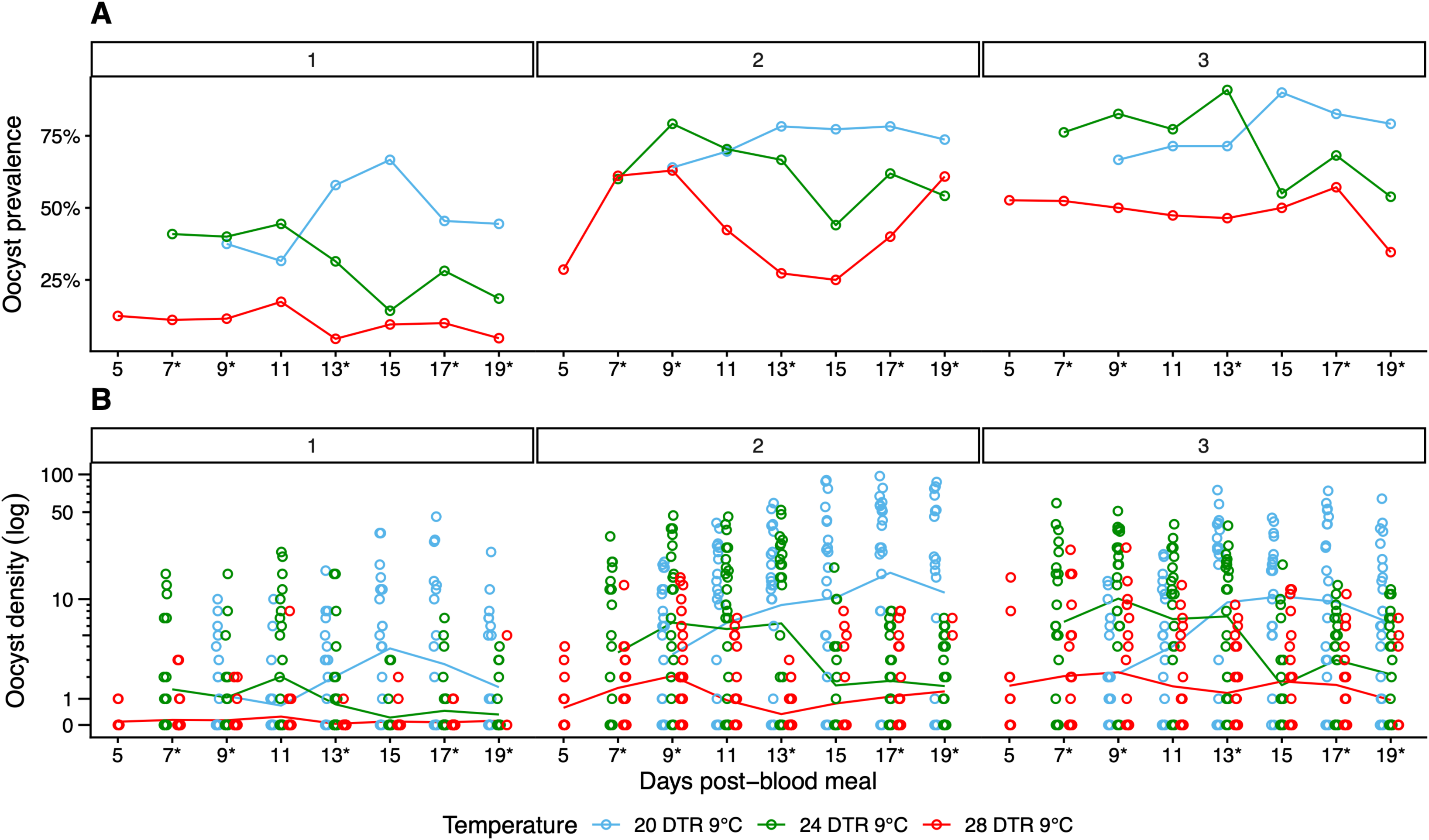
Trends in oocyst prevalence (**A**) and densities (**B**) over time and temperature for each of the three replicates in this study. Lines in (**B**) depict mean, with data points indicating oocyst density in the individual midguts.

**Supplementary figure 2:**
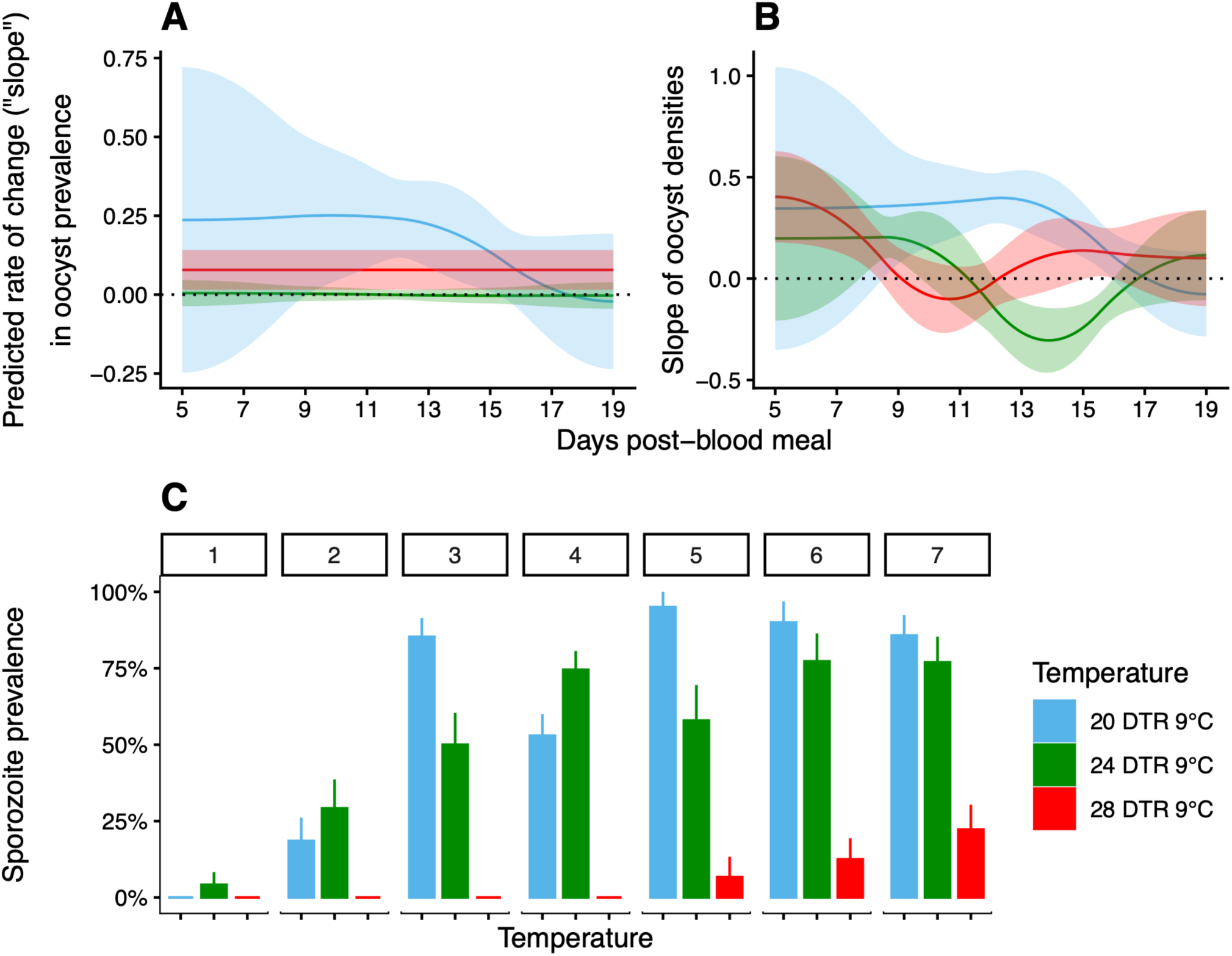
Temperature-specific deviations from the global (‘main effect’) temporal trend in oocyst prevalence (**A**) and density (**B**); sporozoite prevalence in the salivary glands of mosquitoes from each of the seven replicates (**C**). Lines in (**A**) and (**B**) show the effect of temperature on the rate of change, with prevalence expressed as logit/log-odds of infection (**A**) and densities expressed as log- oocyst density per dpbm (**B**); shaded areas represent 95% confidence intervals. In (**C**), replicates 1-7 are ordered in increasing prevalence (averaged over all three temperatures).

**Supplementary figure 3:**
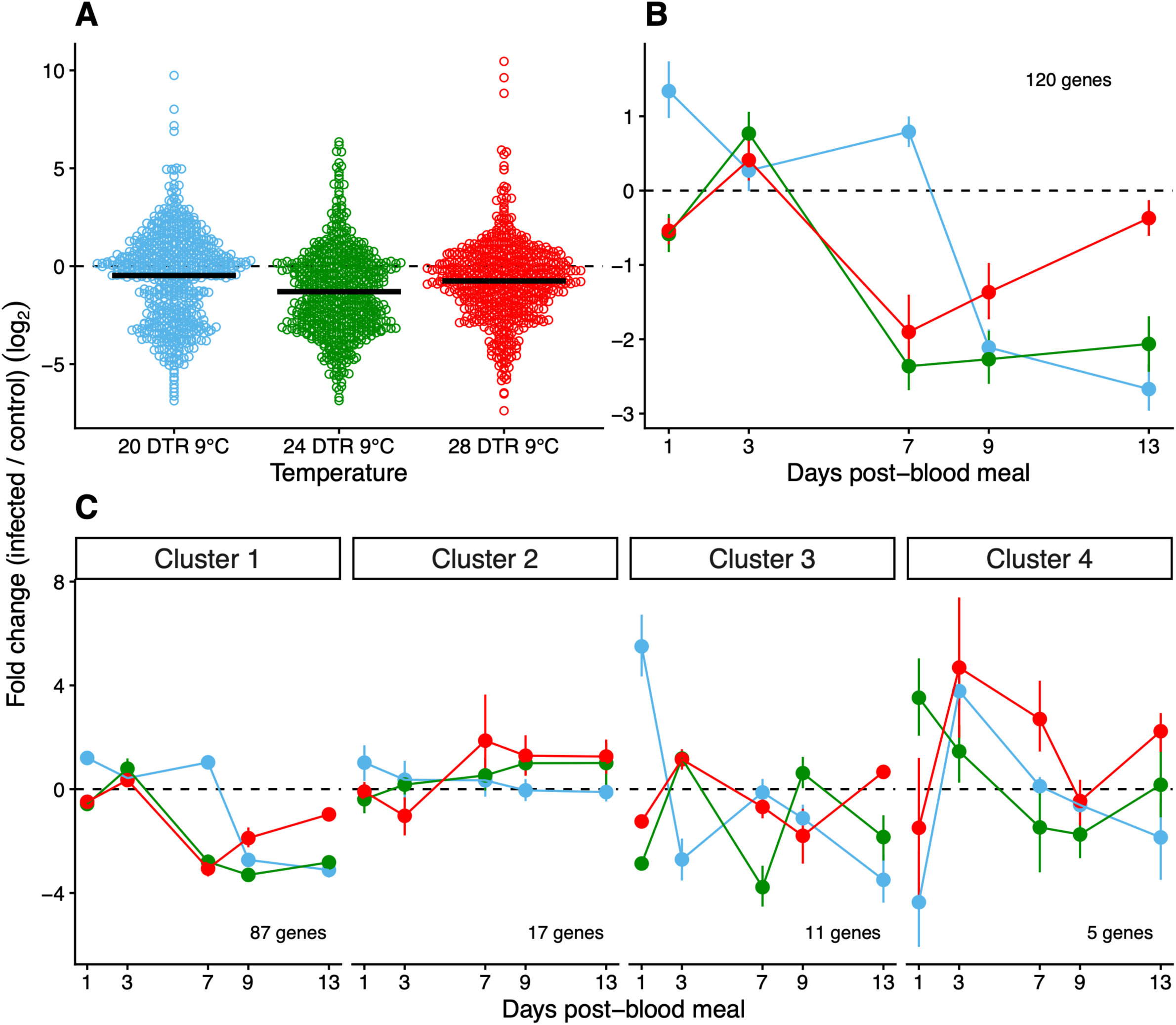
Overall effect of temperature on expression of the 120 genes in infected mosquitoes relative to the controls (**A**), over time and between temperatures (**B**), and separated according to the co-expression clustering; separation was chosen manually after the primary split in the hierarchical dendrogram in Figure 6 (**C**).

**Supplementary figure 4:**
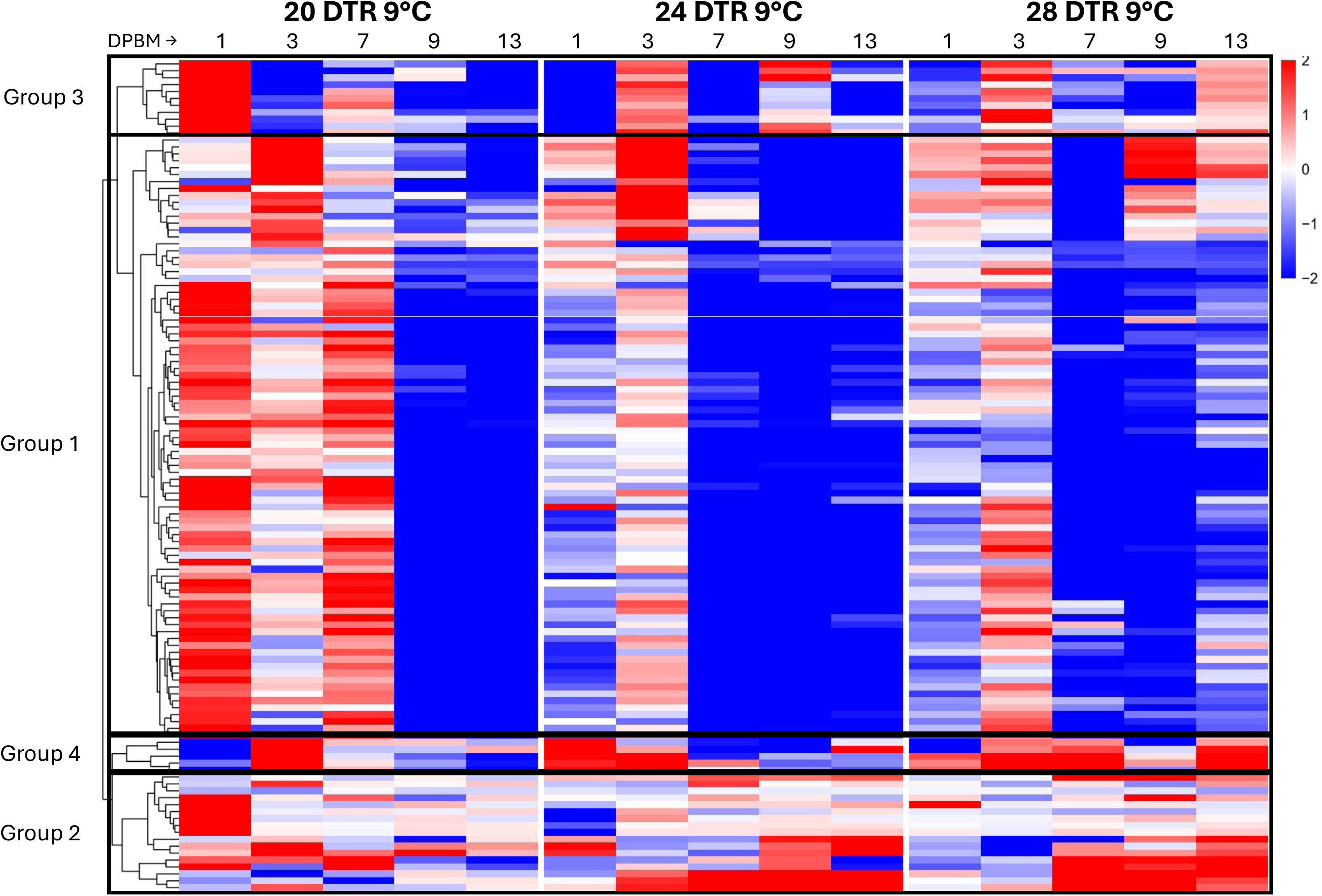
Heatmap incorporating all groups from Figure 6 in the main text. Boxes with black outlines delineate the four groups.

**Supplementary figure 5:**
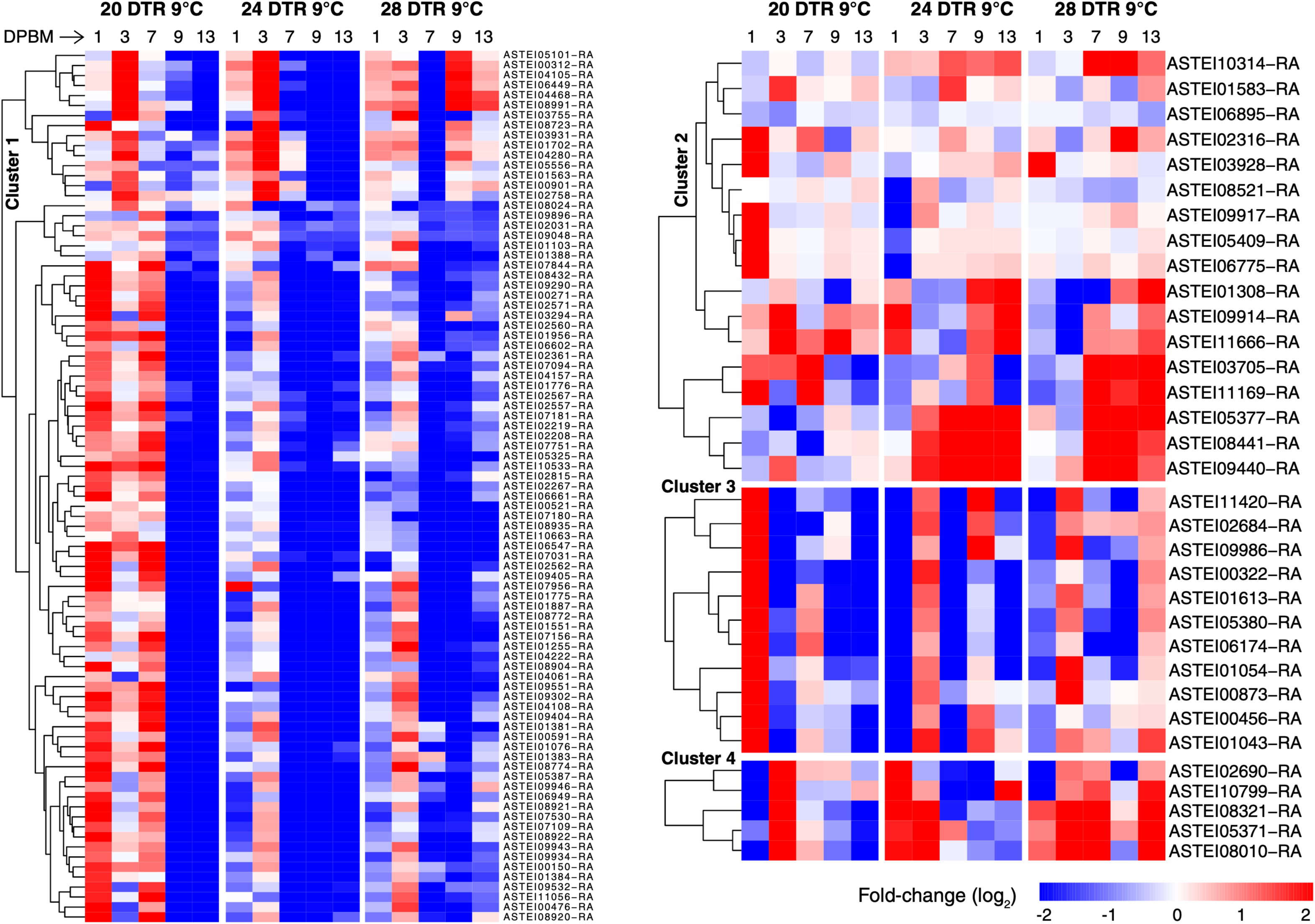
Same as Figure 6 but annotated with VectorBase accession numbers.

**Supplementary figure 6:**
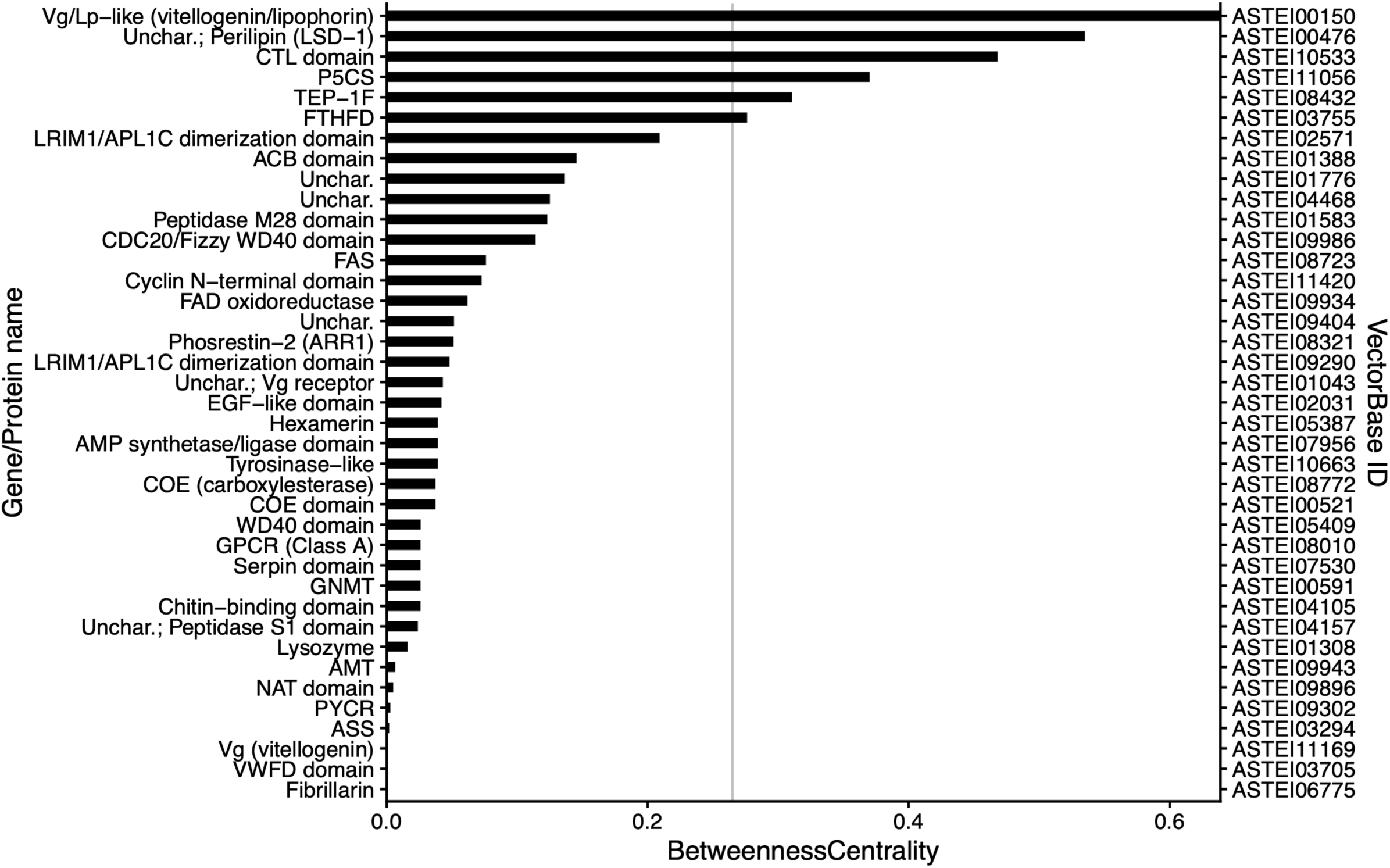
Values of BetweennessCentrality for all proteins with non-zero values from the STRING network; grey line corresponds to x-coordinate of mean + 1SD value of 0.265.

**Supplementary Figure 7:**
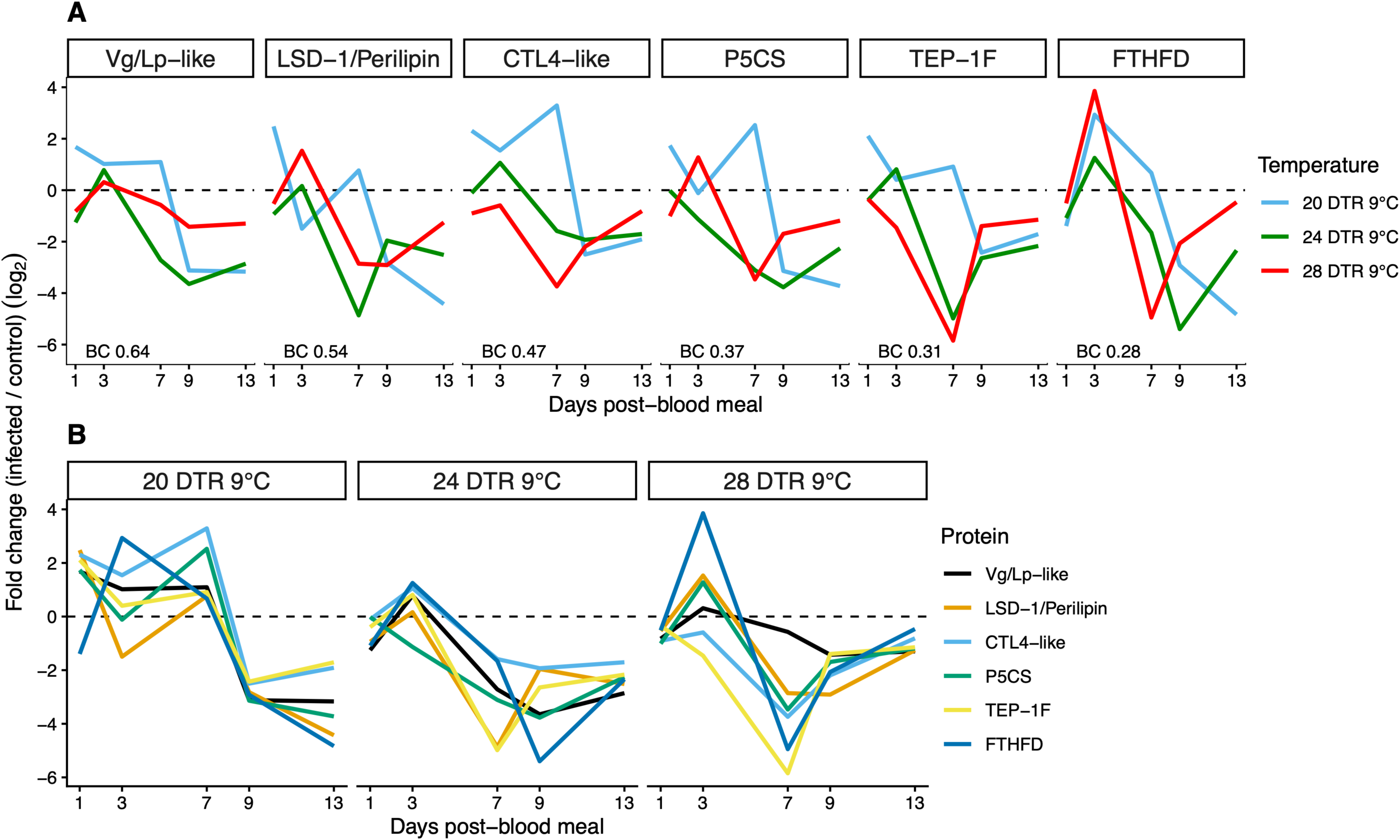
Temporal expression profiles of the six high-centrality proteins (BC > 0.265) across three temperature regimes. Panel (**A**) shows mean log₂ fold change over time, with panes ordered by descending Betweenness Centrality score. Panel (**B**) shows the same data arranged by temperature, with legend and corresponding colors ordered by Betweenness Centrality.

## SUPPLEMENTARY TABLES

**Supplementary Table 1.** Functional enrichment supporting the MCL module labels in Figure 7. Modules were identified from the network in Figure 7 using MCL clustering with a granularity/inflation value of 3. Group-wise enrichment analysis was performed for modules containing ≥4 proteins, and redundant terms were collapsed using a 0.5 cutoff. Figure 7 labels represent manual biological summaries of the retained enrichment terms in the “Description” column. Clustering and enrichment were performed with the STRING database (v12.0) via the stringApp (v2.2.0; released December 2024) implemented in Cytoscape (v3.10.4) ^46^.

| Supplementary table 1 |  |  |  |  |  |
| --- | --- | --- | --- | --- | --- |
| MCL cluster | Category | Term name | Description | # genes (background) | FDR value |
| 1 | SMART Domains | SM00020 | Trypsin-like serine protease | 13 (167) | 1.21E-20 |
| 1 | InterPro Domains | IPR022700 | Proteinase, regulatory CLIP domain | 5 (23) | 5.39E-08 |
| 1 | STRING Clusters | CL:3482 | Peptidase S1A, chymotrypsin family | 6 (66) | 2.78E-07 |
| 1 | KEGG Pathways | map04624 | Toll and Imd signaling pathway | 6 (125) | 7.75E-07 |
| 1 | UniProt Keywords | KW-0720 | Serine protease | 5 (61) | 1.27E-06 |
| 1 | KEGG Pathways | map04080 | Neuroactive ligand-receptor interaction | 5 (248) | 6.20E-04 |
| 1 | Pfam | PF00089 | Trypsin | 3 (22) | 0.0036 |
| 1 | COMPARTMENTS | GOCC:0005576 | Extracellular region | 7 (689) | 0.0081 |
| 1 | STRING Clusters | CL:3603 | Mixed, incl. Intracellular oxygen transport, and Clip or disulphide knot domain | 2 (5) | 0.0208 |
| 2 | UniProt Keywords | KW-0596 | Phosphopantetheine | 2 (5) | 0.0023 |
| 2 | Pfam | PF00372 | Hemocyanin, copper containing domain | 2 (7) | 0.0133 |
| 3 | STRING Clusters | CL:11075 | Mixed, incl. Wnt signaling pathway, and ECM-receptor interaction | 6 (171) | 6.62E-07 |
| 3 | Reactome Pathways | MAP-196854 | Metabolism of vitamins and cofactors | 4 (193) | 0.0058 |
| 3 | STRING Clusters | CL:11400 | Mixed, incl. Egg coat formation, and Regulation of actin filament-based movement | 2 (8) | 0.0138 |
| 4 | STRING Clusters | CL:5942 | Carboxylesterase, type B, and Peptidase M28 | 3 (16) | 1.60E-04 |
| 4 | STRING Clusters | CL:8365 | Ribosome biogenesis, and tRNA modification | 3 (193) | 0.0222 |
| 5 | GO Biological Process | GO:0009084 | Glutamine family amino acid biosynthetic process | 4 (11) | 1.43E-07 |
| 6 | STRING Clusters | CL:2109 | Regulation of APC/C activators between G1/S | 3 (61) | 0.0035 |
| MCL cluster | Category | Term name | Description | # genes (back-ground) | FDR value |
|  |  |  | and early anaphase, and Cyclin-dependent protein kinase holoenzyme complex |  |  |
| 7 | STRING Clusters | CL:9409 | Spliceosomal complex, and Leucine-rich repeat, typical subtype | 5 (158) | 1.09E-06 |
| 8 | KEGG Pathways | map00260 | Glycine, serine and threonine metabolism | 3 (45) | 1.00E-04 |
| 9 | SMART Domains | SM00370 | Leucine-rich repeats, outliers | 3 (131) | 0.0031 |

